# Modular Assembly of Biohybrid Machines Using Force-Enhanced Skeletal Muscle Actuators

**DOI:** 10.1101/2025.07.06.663299

**Authors:** Humphrey Yang, Avery S. Williamson, Ji Min Seok, Daniel Ranke, Duncan McGee, Emily Trotto, Kiyn Chin, Dinesh K. Patel, Adam W. Feinberg, Tzahi Cohen-Karni, Victoria A. Webster-Wood

## Abstract

Muscle-based biohybrid systems integrate living muscle with engineered structures to create soft robots, biological models, and regenerative platforms. However, current actuators often lack strength and are difficult to assemble into complex devices. This study presents a suspended compliant skeleton that enhances muscle maturation by providing passive resistance, enabling high-stroke self-exercise without external stimulation. Using immortalized C2C12 cells, the resulting actuators achieved millimeter-scale strokes and millinewton-scale forces, surpassing previous benchmarks. Magnetic interfaces embedded in the skeleton allowed modular assembly into multi-degree-of-freedom devices such as grippers, arms, and positioning stages. These interfaces also support actuator replacement and repair, improving resilience and scalability. This approach significantly boosts engineered muscle performance and offers a robust, modular platform for building high-functioning, repairable biohybrid machines.

---

Across the natural world, animals demonstrate levels of adaptability, robustness, and performance that continue to inspire, and often elude, robotics research. The multifunctional, gentle dexterity and strength found in living systems, from human hands (*1*) to elephant trunks (*2*), continues to drive innovation in the field. However, a key gap between animals and modern robots remains with respect to the actuators, with muscles exhibiting adaptability (*3*), high power density (*4, 5*), and the ability to self-heal (*6*) and self-assemble (*7, 8*), while traditional robotic actuators require bulky housings and support infrastructure (*9, 10*), high voltage (*4, 11*), or high temperature (*4, 12*) to drive actuation. Although some traditional robotic actuators outperform muscle in individual performance metrics, muscle meets key engineering needs across multiple dimensions of performance in ways that synthetic alternatives do not (*3, 13, 14*).

The emerging field of biohybrid robotics seeks to address this gap by directly harnessing biological muscles as actuators (*15–20*). In this paradigm, muscles are often integrated with an abiotic structure tailored to specific needs and use cases. The muscle then serves as an actuator to drive the device in response to external stimulation. Using a biohybrid approach, researchers have created devices capable of crawling (*21, 22*), swimming (*23, 24*), pumping (*25, 26*), and gripping (*22, 27*). Beyond these example devices, biohybrid actuators and machines have broader applications, including the creation of lab-on-chip systems for biological (*24*) or medical studies (*28*) and sustainable robotic systems (*29*). The muscle actuators used in biohybrid machines typically originate from either explanted tissue (*27, 30*) or are created using *in vitro* culture techniques from tissue engineering (*21–26*). The former often produces more force but has limited availability due to the requirement of harvesting from living animals and limits the shape and scale of the actuator to that produced by the source animal. In contrast, *in vitro* culture approaches are customizable, and actuators can be fabricated on demand (*31*). However, the maturation and assembly of muscle actuators remains an ongoing barrier to the broader adoption of cultured biohybrid devices in robotics.

In the manufacture of cultured biohybrid devices, while there are established methods for the initial fabrication of muscle-based actuators, such as casting (*32–34*) and bioprinting (*31, 35, 36*), providing adequate developmental conditions and constructing complex multiactuator devices remains a bottleneck. Due to their fragility and softness, cultured muscle actuators are typically manufactured directly onto the target device structure (*21, 34, 37, 38*) to minimize handling. Alternatively, muscle actuators can be fabricated and matured on a scaffold before being manually detached and transferred to the target device (*39, 40*). The latter method may enable modular biohybrid device construction, but over-handling of muscle actuators may compromise their structural integrity. Although the former approach mitigates handling needs and risk, it presents a challenge in building devices that require multiple muscle actuators, as a defective actuator could render the entire device non-functional. Furthermore, the mechanical properties provided by the culture environment throughout maturation are crucial as they directly influence cell alignment and maturity (*39, 41, 42*). As a consequence, there is a need for manufacturing approaches that allow the flexibility to differentiate and mature muscle actuators on engineered scaffolds, providing the appropriate developmental cues, while being able to safely and robustly transfer and assemble the actuators onto more complex robotic devices for application.

In addition to challenges in manufacturing with living muscle, the design space of biohybrid robots is limited by the current muscle actuator’s force capabilities. Indeed, the phenotype of most engineered muscle actuators remains relatively immature, with force capabilities falling short of native muscle tissue (*15*). To address this, researchers have found that the performance of these actuators can be increased by exercising the muscle during maturation (*31, 39, 43, 44*). This exercise can be induced through external stimulation by applying an electric field to the culture (*37, 45*) or optogenetically by modifying the cells to contract in response to regularly pulsing light in the culture environment (*32, 46–48*). However, these approaches require bulky stimulation equipment or modifications to the cells. While such methods improve actuator force capabilities in isolation, the matured muscle actuators performance still fall short of native muscle (*15, 17*). On the other hand, engineering compliant skeletons to provide passive stimulation has emerged as a simple yet effective method for culturing biohybrid devices that can self-exercise (*39–41*). This approach leverages the springiness of compliant mechanisms to provide passive resistance against the spontaneous contraction of muscles, thus eliminating the need for an external stimulation apparatus. However, engineering compliant skeletons to provide proper maturation cues and support their modular construction for application on devices remains an open challenge.

Building upon compliant mechanism literature, this work presents a method for culturing high-performance muscle-based biohybrid actuators and modularizing the construction of biohybrid devices. To promote the linear contraction development of the muscle-based actuator, we have designed compliant skeletons with tailored single degree-of-freedom (DOF) mobility (Fig. 1 A). Taking inspiration from how animals gain muscle strength from repeated workouts, we also engineer the skeletons to facilitate large-stroke exercise and provide resistance against spontaneous contractions during maturation. The resulting actuators are capable of producing mm-scale strokes (up to 10.56% of actuator rest length) and mN-scale forces (up to 2.09 mN) under electrical stimulation (Fig. 1B), thereby competitively navigating a unique performance space (*32–34, 38, 39, 49–56*) in cultured biohybrid robotics (Fig. 1C). In particular, the presented muscle-based biohybrid actuators achieve similar metrics as harvested murine skeletal muscle cells (*34*), while being fabricated from the immortalized C2C12 cell line. The compliant skeletons also feature magnetic interfaces, allowing matured actuators to be assembled into the structure of a target device (Fig. 1D). During the process, the compliant skeleton provides structural support to the muscle actuator, reducing the risk of damage during device handling. Once installed, the skeleton can be cut away to release the actuator. The interface also enables device repair and reconstruction following damage and partial actuator loss, making biohybrid devices more economical and resilient.

**Figure 1:**
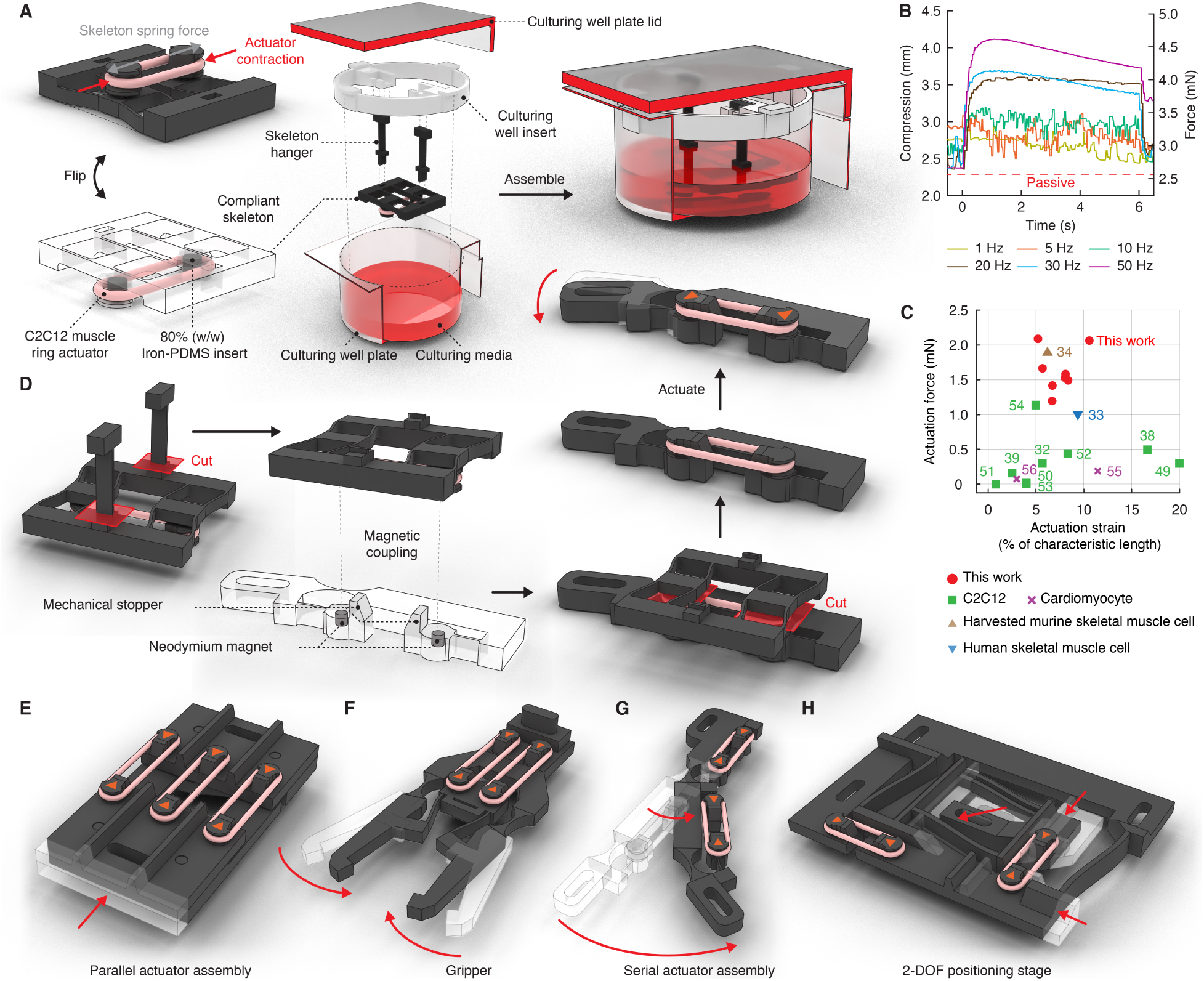
Muscle-based biohybrid actuator maturation and modular assembly through compliant skeletons and magnetic interfacesA) The suspended self-stimulating compliant skeleton and culturing setup. B) A representative electrical stimulation response of a 20 mm long actuator matured on a 1.121 mN/mm skeleton. C) Performance space comparison between this work and prior literature (*32–34, 38, 39, 49–56*). D) Schematic depiction of the assembly of matured muscle actuators into an external, compliant structure to create a biohybrid device. E) A multi-fiber linear actuator assembled from a compliant structure and three parallelly arranged muscle actuators. F) A compliant gripper driven by two parallelly arranged muscle actuators. G) A compliant robotic arm driven by two serially arranged muscle actuators. H) A 2-DOF positioning stage driven by two orthogonal actuators, each controlling the displacement of one axis.

The modularity and actuation capability of the proposed approach enabled the construction of complex devices in the mesoscale. Several devices were implemented to illustrate different assembly strategies. In a parallel assembly (Fig. 1E), the actuators can provide larger force output and redundancy, ideal for making devices on a larger scale that require high driving force, such as a 50 mm scale manipulator (Fig. 1F). When arranged in series (Fig. 1G), this approach supports devices having an amplified motion range and alignment of actuators in orthogonal orientations enables the generation of motions along multiple degrees of freedom (Fig. 1H). Together, the compliant mechanism-based maturation methods and modular magnetic assembly presented in this work greatly enhance the force capabilities of biohybrid actuators and the possible design space of biohybrid robotics.

## Results

### Passive compression of compliant skeletons

Compliant skeletons that accommodate varying actuator lengths (Fig. 2A) with a targeted translational degree of freedom (Fig. 2B) were fabricated (Fig. 2C). The skeletons had a linear load curve within the intended actuator compression range (Fig. 2D, Fig. 2E). Two skeleton stiffness designs were explored: equivalent stiffness (Fig. 2F) or equivalent strain (Fig. 2G). Over the maturation period in differentiation media (DM), the skeleton exhibited increasing passive compression resulting from the muscle actuator’s tissue compaction (Fig. 2H). All skeleton groups exhibited the same degree of passive compression on day 0 (D0), immediately after transfer from the cast wells to the compliant skeletons. The amount of passive compression subsequently increased with time (Fig. 2I). In the equivalent stiffness skeleton group, the development was relatively linear between D0 and D14. In contrast, in the equivalent strain skeleton group, a rapid increase in the skeleton’s passive compression was observed starting from D2, but eventually plateaued at around D10. On D14, the amount of passive compression largely depended on the skeleton’s stiffness. Across the equivalent stiffness groups, there was no statistically significant difference in the amount of passive compression between actuators of different lengths (Fig. 2J). Conversely, there was a statistically significant difference in passive compression across skeletons of varying stiffness. Given the same actuator length, the softer skeletons were significantly more compressed than their stiffer counterparts (*p* < 0.05). The differences were also proportional to the stiffness of the skeleton. Within the equivalent strain skeleton group, the compression amount was also proportional to the length of the actuator. Despite these differences, when inspecting the passive compression forces exerted by muscle actuators cultured on different skeletons (Fig. 2K), statistically significant differences were only observed between the 15 mm and 10 mm skeletons in the equivalent stiffness group (*p* < 0.05).

**Figure 2:**
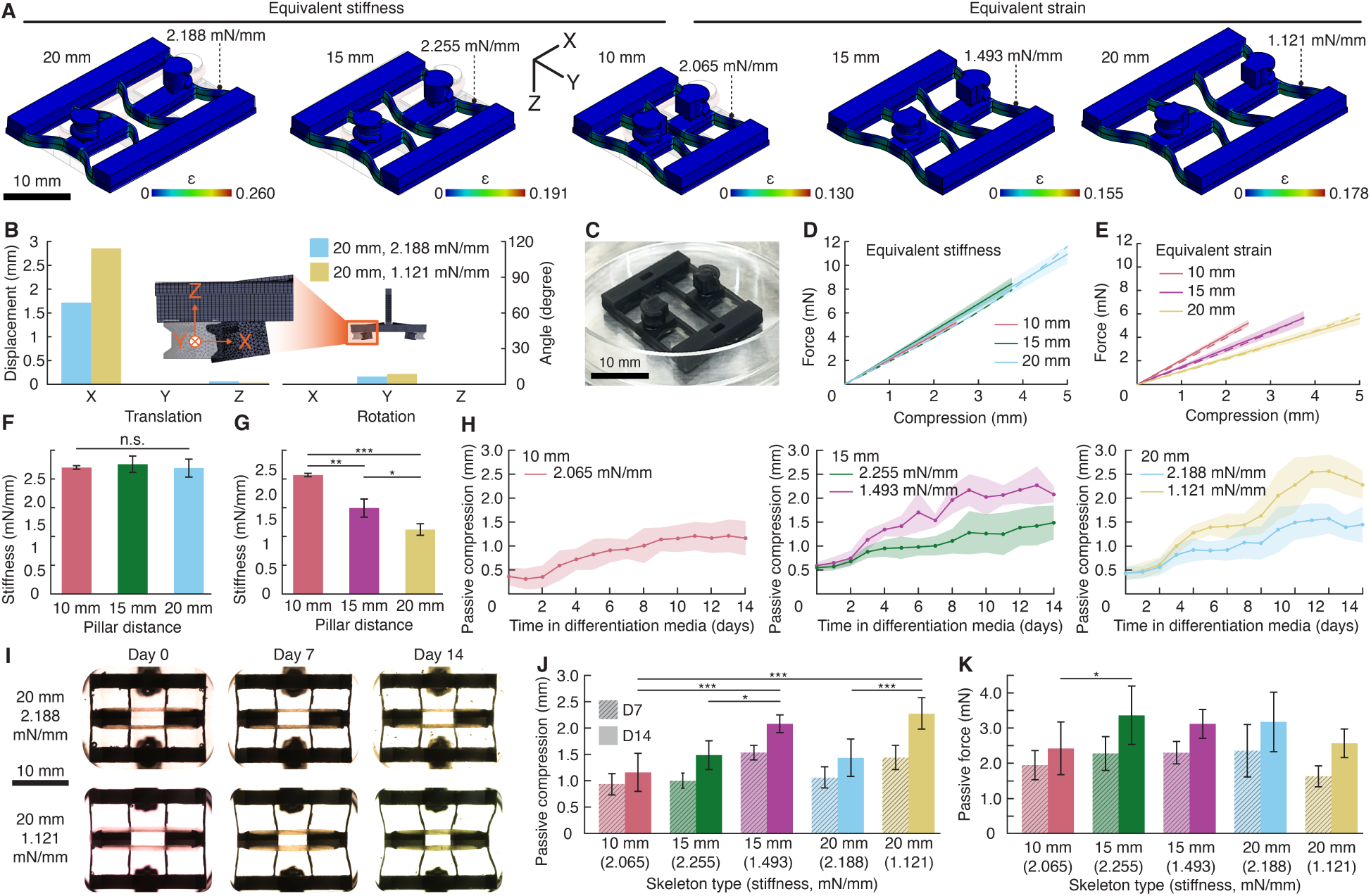
The design and characterization of compliant skeletons, and maturation of muscle actuators. A) Simulations of skeletons under 25% strain (from left to right: 10, 15, and 20 mm). The color spectrum shows the Von Mises strain. Scale bar, 10 mm. B) Simulated skeleton DOF analysis in response to a 6 mN compression force. Inset, simulated skeleton deformation. C) Fabricated 10 mm compliant skeleton before mounting the muscle actuator. Scale bar, 10 mm. D) Skeleton mechanical property characterization of the equivalent stiffness group. Solid lines, physical experiment, N=3; dashed lines, simulation. E) Skeleton mechanical property characterization of the equivalent strain group. Solid lines, physical experiment, N=3; dashed lines, simulation. F) Skeleton stiffness comparison within the equivalent stiffness group. N=3. G) Skeleton stiffness comparison within the equivalent strain group. N=3. H) Passive skeleton compression over time. N=5 to 7. I) Representative images of 20 mm skeletons on D0, D7, and D14. Scale bar, 10 mm. J) Passive skeleton compression versus skeleton designs by length and (stiffness). N=5 to 7. K) Passive actuator force versus skeleton designs by length and (stiffness). N=5 to 7. All data are mean±SD. Statistically significant difference threshold for Tukey’s post hoc test: *, 0.01 ≤ *p* < 0.05; **, 0.001 ≤ *p* < 0.01; ***, *p* < 0.001.

### Spontaneous self-exercise dynamics

Over the maturation period, the actuators’ spontaneous contractions exhibited a shift in their prominent frequency and amplitude over time (Fig. 3A). On D7, spontaneous activities were dominated by high-frequency, low-amplitude contractions. The contractions then reduced in frequency but grew in amplitude over time. In the 10 mm and 15 mm actuator length groups, the shift in spontaneous contraction frequency and amplitude persisted from D7 to D14. Figure 3B shows the representative spontaneous contraction patterns of 10 mm actuators (see also Video S1). In the spectral domain (Fig. 3C), the prominent contraction frequency halved from D7 to D10 and was reduced by an order of magnitude from D7 to D14. The maximum stroke of spontaneous contractions also increased with maturation time (Fig. 3D). In contrast, in the 20 mm actuator groups, the maximum stroke of spontaneous contractions peaked on D10. We speculate that the reduction in spontaneous contraction may be caused by nutrient depletion and waste accumulation in the culture medium despite daily media change, as can be seen in Fig. 2I, where the change in the color of the medium suggests acidic waste accumulation. Future work may investigate this effect and further optimize the media change frequency accordingly.

**Figure 3:**
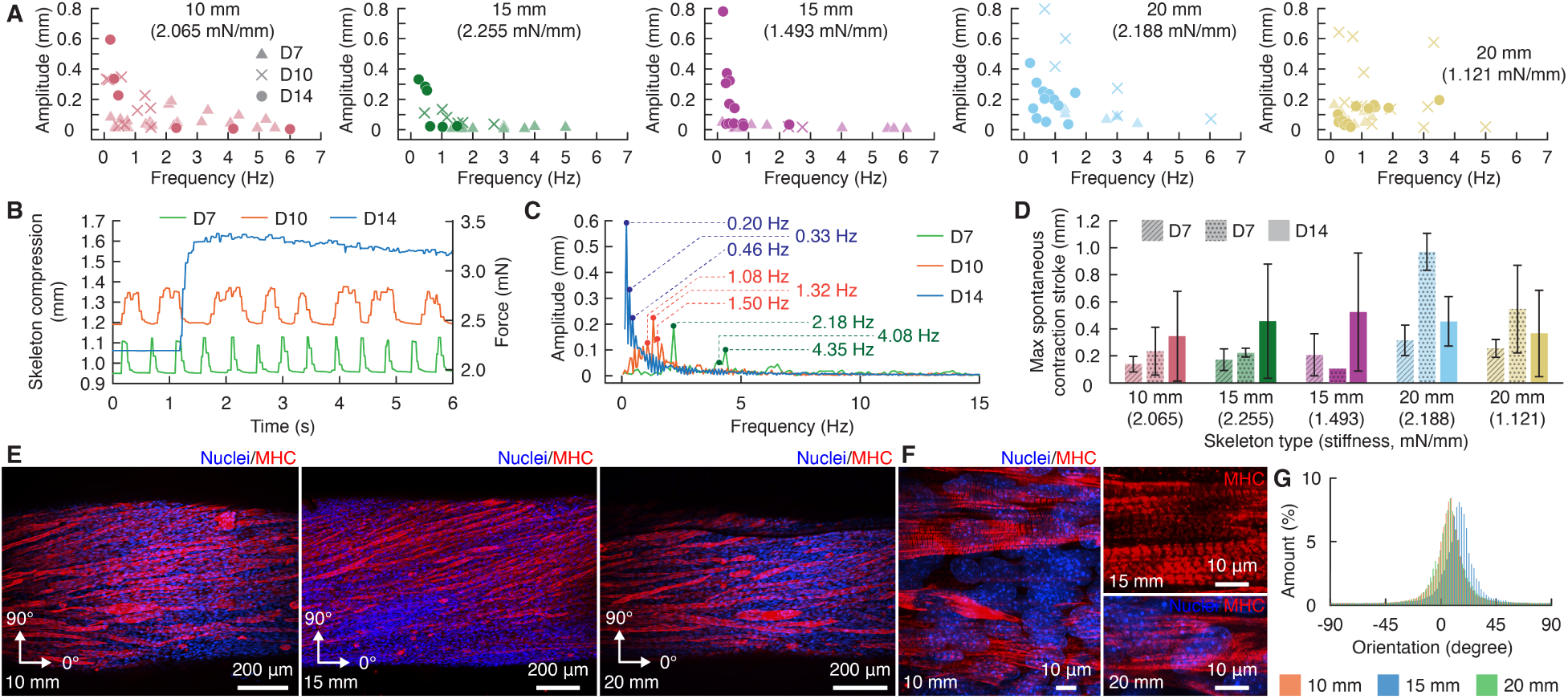
Muscle actuator spontaneous contraction analysis and immunohistochemistry. A) Plots of the three most prominent spontaneous contraction patterns for each actuator type on D7, D10, and D14. N=1 to 8. B) Representative spontaneous contraction patterns of the 10 mm actuator on D7, D10, and D14. C) Fast Fourier transformation results of the patterns presented in (B) with the three most prominent frequencies highlighted. D) Plot of the maximum spontaneous contraction stroke of muscle actuators on D7, D10, and D14. N = 1 to 8. Statistical analysis was omitted due to inconsistent sample sizes. E) Immunohistochemistry of equivalent stiffness actuators on D14. Blue, nuclei; red, myosin heavy chain. Scale bars, 200 *μ*m. F) High-magnification immunohistochemistry of equivalent stiffness actuators on D14 showing striation along myotubes. Blue, nuclei; red, myosin heavy chain. Scale bars, 10 *μ*m. or 10 *μ*m. The arrow indicates striation. G) Myotube alignment histogram of (E). All data are mean±SD. Statistically significant difference threshold for Tukey’s post hoc test: *, 0.01 ≤ *p* < 0.05; **, 0.001 ≤ *p* < 0.01.

### Myotubes align along actuator axis

Immunohistochemistry of the muscle rings confirmed myotube formation along the actuator (Fig. 3E) as well as striations within the myotubes (Fig. 3F). The myotubes were oriented along the actuator’s longitudinal axis with a slight slant (Fig. 3G), similar to that observed in two-dimensional C2C12 culture (*57*).

### Biohybrid actuator force and stroke

The biohybrid actuators cultured on suspended compliant skeletons exhibited active force production in the mN scale and active stroke in the mm scale when subjected to electrical stimulation (Fig. 4A). We characterized all actuators with six-second bursts of stimulation at frequencies ranging from 1-30 Hz, and additionally tested the 15 and 20 mm actuators in the equivalent strain group at 50 Hz stimulation. As expected for biological muscle, the frequency of stimulation resulted in different active stroke responses (Fig. 1B, Fig. 4B, Video S2). At 1 Hz, while the actuators were reacting to the stimulation pulses, spontaneous contractions were also observed between pulses. At 5 Hz, the pulse stimulation led to rapid oscillation in skeleton compression. At 10 Hz, a tetanus contraction plateau started to emerge, but the skeleton compression readout still included higher-frequency oscillations, likely due to spontaneous activity. The oscillation was attenuated at 20 Hz, where the tetanus plateau became apparent and stable across the stimulation window. At higher frequencies (30 Hz and 50 Hz), the tetanus plateau was distinct but also showed signs of relaxation over the stimulation window, indicating that the actuators could no longer sustain their maximum force output after the initial contraction. However, over repeated stimulation (5-10 times) at 30 or 50 Hz, the actuators did not exhibit noticeable long-term fatigue effects or declining performance, suggesting that they could be repeatedly operated at high stimulation frequency without compromising maximum force output. It is worth noting that in this frequency range, the actuators were observed to relax into either spontaneous contraction or complete relaxation after stimulation, and the actuators’ response to stimulation appeared most prominent in episodes where spontaneous activity was not observed between stimulation windows.

**Figure 4:**
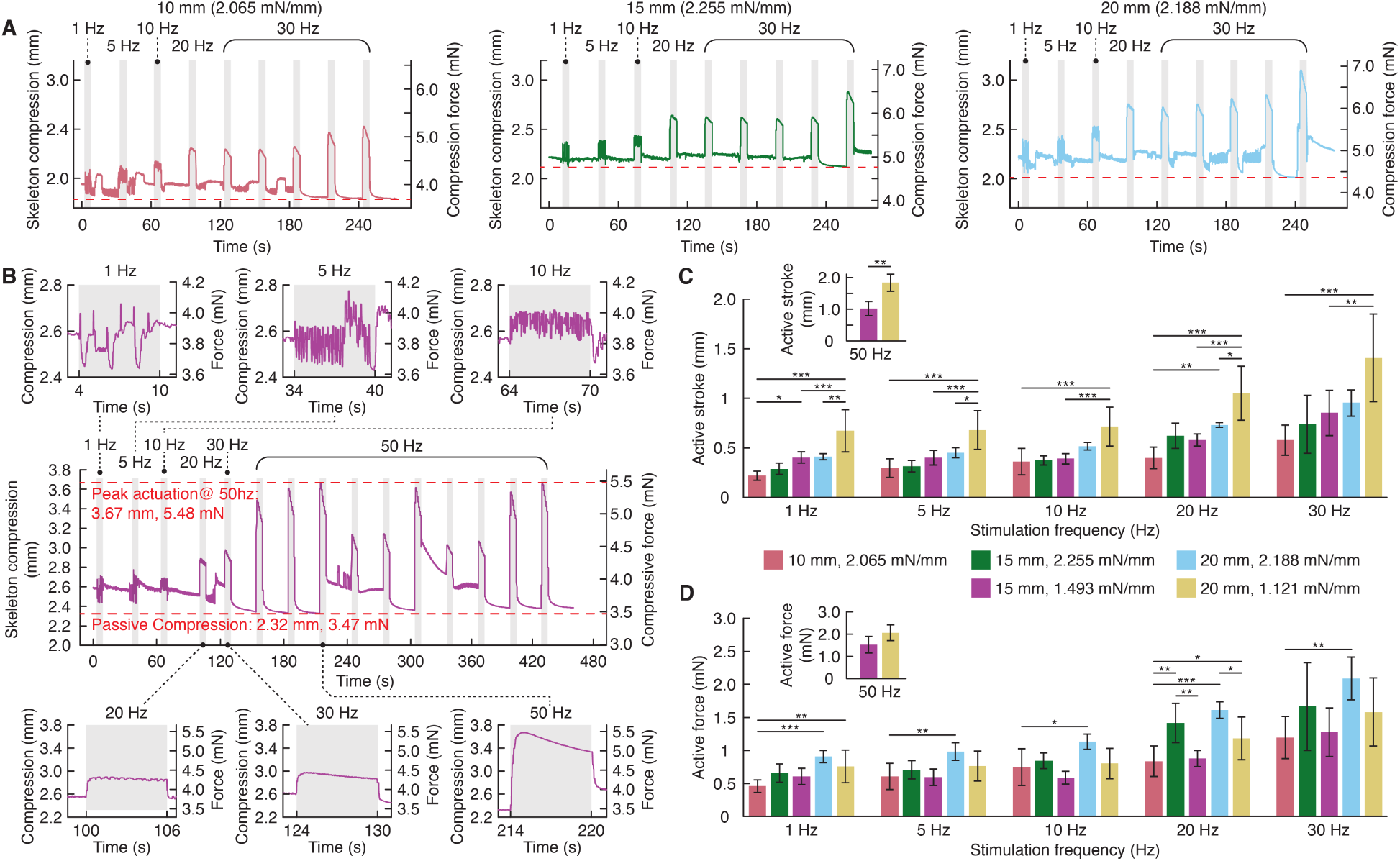
Muscle Actuator active stimulation analysis. A) Representative stimulation response of muscle actuators matured on the equivalent stiffness skeletons. The dashed red line shows the baseline passive compression. B) Representative electrical stimulation response of a 15 mm actuator cultured on the equivalent strain skeleton. C) Actuator active stroke at varying stimulation frequencies. N=4 to 10 (insets: N=3 to 4). D) Actuator active force production at varying stimulation frequencies. N=4 to 10 (insets: N=3 to 4). All data are mean±SD. Statistically significant difference threshold for Tukey’s post hoc test: *, 0.01 ≤ *p* < 0.05; **, 0.001 ≤ *p* < 0.01; ***, *p* < 0.001.

Under the same stimulation frequency, the degree of simulation response differed by the skeleton design, with longer and softer skeletons leading to higher active stroke (Fig. 4C, Video S3). The largest active stroke was observed in actuators cultured on the 20 mm, equivalent strain skeleton, reaching 1.41 ± 0.44 mm at 30 Hz and 1.84 ± 0.27 mm at 50 Hz. In the equivalent stiffness skeleton group, actuators cultured on different skeletons exhibited a similar range of active stroke at higher stimulation frequencies. Conversely, in the equivalent strain skeleton group, the 20 mm actuator had significantly higher active stroke than the shorter actuators (10 mm actuator: 0.58±0.15 mm; 15 mm actuator: 0.85±0.23 mm)at a 30 Hz stimulation frequency (*p* < 0.01). When factoring in skeleton stiffness, the muscle actuators had a similar force output regardless of the skeleton they were matured on (Fig. 4D). However, the 20 mm actuators, particularly those in the equivalent stiffness group, consistently and significantly produced more force than the 10 mm actuators (*p* < 0.05), suggesting that a stiffer skeleton and longer actuator may result in better performance.

### Multi-actuator biohybrid devices through modular assembly

The actuators’ modularity, enabled by the magnetic interface, allowed assembly into different arrangements post-maturation. In our device demonstrations, the target device structures were implemented as monolithic compliant mechanisms using the same material that constituted the compliant skeleton, thereby eliminating the need for traditional hinges and bearings. When assembled in a parallel arrangement (Fig. 1E), an array of actuators can provide larger force output and redundancy against partial failure. To demonstrate this ability, three muscle actuators cultured on the 20 mm, 1.121 mN/mm skeleton were assembled in parallel to create a multi-fiber linear actuator (Fig. 5A, Video S4). The amount of passive compression at the muscle actuators’ rest state (Fig. 5B) and active stroke (Fig. 5C) increased with additional actuators, but the relation was not linear. Specifically, while an individual muscle actuator in the triple or double actuator setting generated a similar amount of force, the single actuator setting produced significantly (*p* < 0.05) more force per actuator (Fig. 5D). We speculate that this is likely due to the change in skeleton resistance leading the muscle actuator to rest at a different passive compression length, changing their strain pre-load and causing the actuators to respond in different force production regimes of their length-tension curves. Nonetheless, the results suggest that in a parallel setting, multi-fiber linear biohybrid actuators can retain partial function even when partially damaged. When left with just one actuator, the device was still able to produce approximately half of the force as three actuators (Fig. 5C). Due to the modular nature of our actuators and the separation of maturation and device assembly, the damage could also be repaired by replacing the damaged units to recover the device’s performance (Fig. 5E), thus enabling biohybrid systems to be more resilient and economical.

**Figure 5:**
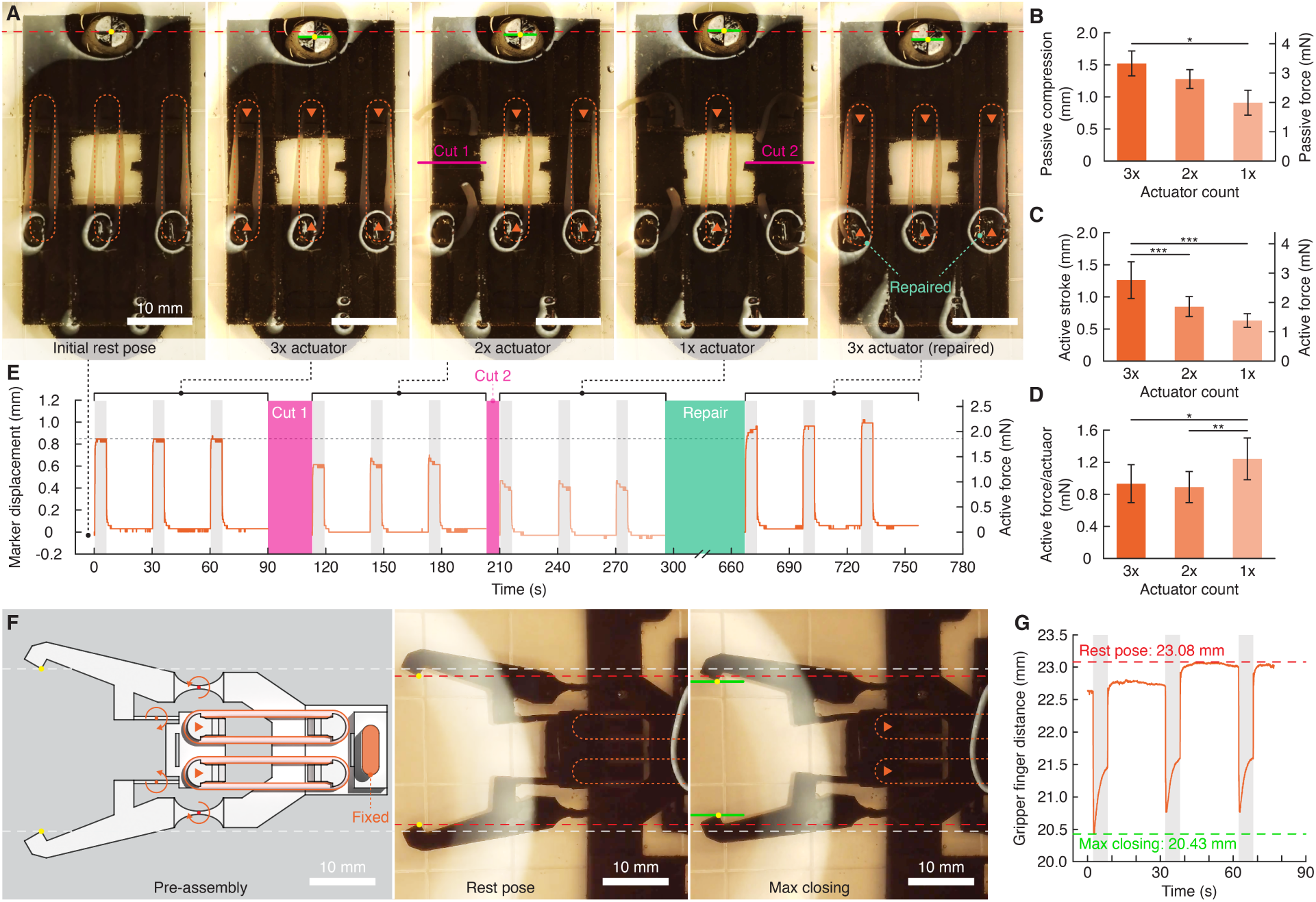
Modular assembly of muscle actuators in parallel arrangements. A) A multi-fiber biohybrid linear actuation device driven by three modular muscle actuators. Scale bars, 10 mm. B) The end-to-end compression of the linear actuation device with different numbers of modular actuators attached. N=9. C) Linear actuation device stroke and force output as a function of the number of modular actuators attached. N=9. D) Linear actuation device force production per actuator with varying numbers of functional modular actuators. N=9. E) An exemplary sequence of the linear actuation device’s performance change due to damage (manual cuts) and repair to individual modular actuators. The gray bands show electrical stimulation windows. F) The design and operation of a compliant gripper. Scale bars, 10 mm. G) The finger distance change of the compliant gripper in response to electrical stimulation. The gray bands show electrical stimulation windows. All data are mean±SD. Statistically significant difference threshold for Tukey’s post hoc test: *, 0.01 ≤ *p* < 0.05; **, 0.001 ≤ *p* < 0.01; ***, *p* < 0.001.

A parallel actuator assembly could also be useful in driving larger-scale biohybrid systems. A biohybrid compliant gripper (Fig. 1F) was fabricated with twice the characteristic length of that demonstrated by the literature (*49*), thus having higher load requirements. Unlike literature that mounted grippers vertically to avoid the device from bending under gravity, we opted to thicken the compliant hinges to withstand the device’s own weight, allowing the gripper to be mounted horizontally (Fig. 5F) at the cost of increased actuation stiffness. Using two parallel actuators, the device was capable of 2.65 mm of closing under electrical stimulation (Fig. 5G, Video S5).

Assembling actuators in series enables a device to have a larger work space (Fig. 1G). To demonstrate this ability, a robotic arm was designed and assembled to leverage two serially arranged muscle actuators (Fig. 6A, Video S6). Each actuator drives a rotational elbow joint of identical stiffness, amplifying the end effector’s rotational range of motion (Fig. 6B). Unlike the parallel actuator arrangement, the range of motion was roughly proportional to the number of actuators in the serial system. When partially damaged, the arm’s angular workspace (i.e., rest pose and active stroke) was halved by the malfunctioning actuator (Fig. 6C). The compromised workspace could then be restored by replacing the compromised actuator, leveraging the modular magnetic interfaces.

**Figure 6:**
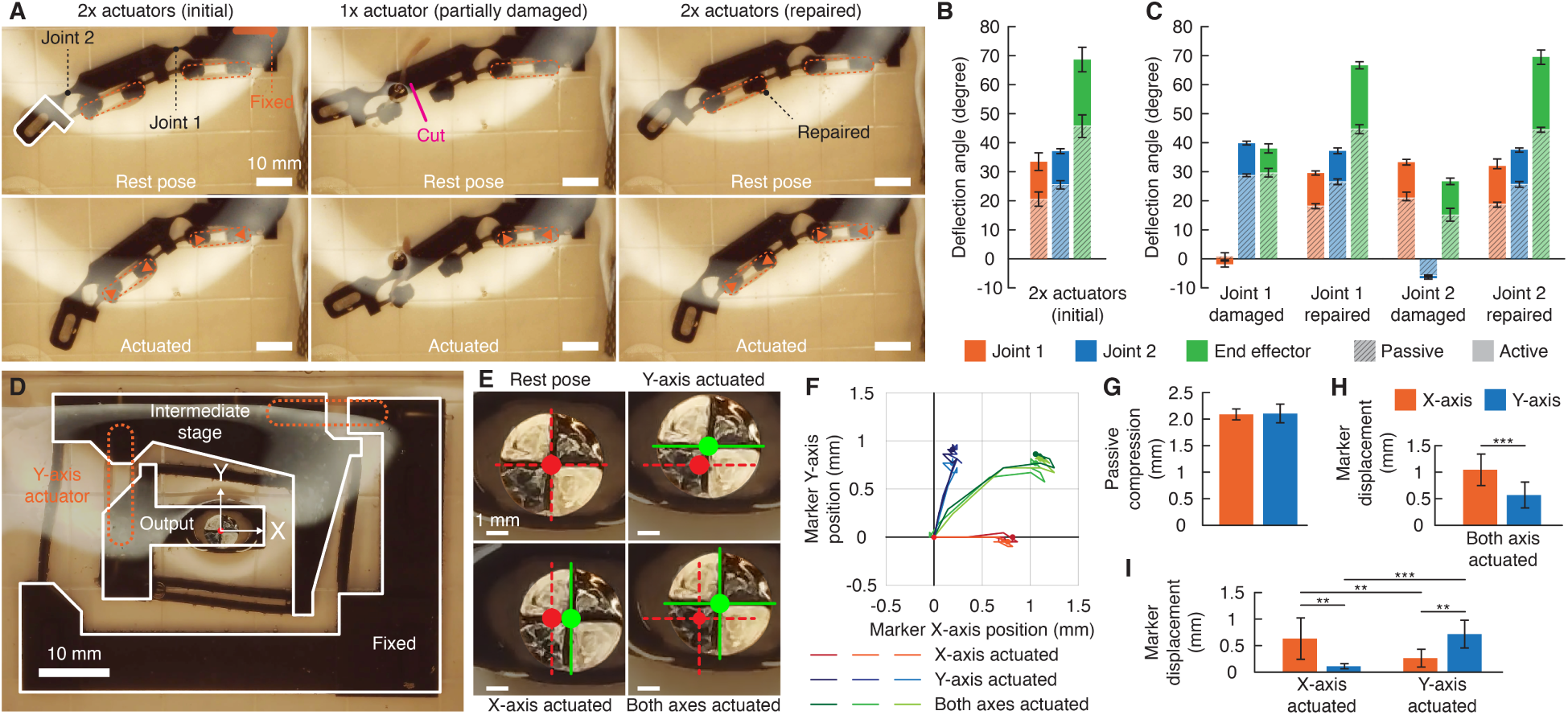
Modular assembly of muscle actuators in serial arrangements. A) A robotic arm-like device comprising two muscle actuators. Scale bars, 10 mm. B) Plot of the arm-like device’s passive and active angular displacement. N=9. C) Plot of the arm-like device’s motion range after damage and repair. N=3. D) A 2-DOF motion stage comprising two muscle actuators driving the X- and Y-axis displacement. Scale bars, 10 mm. E) Images of the 2-DOF motion stage output tracker position change under different actuation modes. Red mark, rest post; green mark, actuated. Scale bars, 1 mm. F) The output tracker trajectory of the 2-DOF positioning stage under different actuation modes. N=3. G) The 2-DOF motion stage’s passive compression along each axis. N=3. H) The 2-DOF motion stage’s output tracker displacement under simultaneous X- and Y-axis stimulation. N=9. I) The 2-DOF motion stage’s output tracker displacement under targeted axis actuation. N=9. Subfigures G-I share the same legend. All data are mean±SD. Statistically significant difference threshold for Tukey’s post hoc test: **, 0.001 ≤ *p* < 0.01; ***, *p* < 0.001.

Individual actuators can also be designed to drive distinct DOFs in a kinematic system, creating a multi-DOF positioning device (Fig. 1H). To demonstrate this, we implemented a 2-DOF motion stage with two orthogonal muscle actuators displacing the output stage along the X- and Y-axis (Fig. 6D, Video S7). By positioning the electrodes for selective stimulation, the output stage could be individually or jointly displaced along the two axes (Fig. 6E, Fig. 6F). Noticeably, while each axis used the same actuator type and was designed to have identical stiffness, leading to similar passive compression in response to the muscle actuator’s compaction tension (Fig. 6G), there was a significant difference in active stroke when simultaneously actuating both axes (Fig. 6H, *p* < 0.001). This is likely caused by the relative angle between the actuators and the stimulating electric field, which was aligned to the X-axis. When stimulating individual degrees of freedom, while the targeted axis displayed significantly higher displacement than the non-stimulated axis (*p* < 0.01), there was cross-talk between actuators, causing a fractional displacement along the other axis. Future research may consider fine-tuning the stimulation condition to mitigate this effect or exploring integrated electrodes for improved stimulation specificity (Fig. 6I).

## Discussion

The results presented here suggest opportunities to further enhance the performance of biohybrid muscle-based actuators. While the actuators demonstrate high force and stroke production, their performance is still not comparable to skeletal muscles in animals, which often have a 20% fractional shortening. Optimizing the fabrication method, such as employing 3D printing to avoid casting and manual muscle ring transfer and create fascicle-like structures (*36,58*), more aggressive stimulation conditions (*49*), modified bioink formulation (e.g., using human (*33*) or chick (*59, 60*) skeletal muscle cells), and compliant skeleton design, could also strengthen actuator performance. In particular, taking inspiration from biological skeletal and muscular structure, culturing muscle actuators as antagonistic pairs on a compliant skeleton may further intensify the mechanical stimulation they receive, thus enhancing their functional development. Furthermore, constructing the compliant skeletons with stiffness-changing materials (*61, 62*) may also allow the skeleton to adapt its stiffness to growing actuator strength, providing dynamic and progressive resistance for the actuator’s maturation.

On the other hand, this work also demonstrates some of the challenges and opportunities in biohybrid device design. As has been consistently observed in the field, actuator scale and force remain limited due to the lack of vascularization within these tissues. Future integration of vascularization or other means of oxygenation will play a critical role in scaling up muscle-based actuators or broader use in robotics and translational medicine. As shown in the device examples, there is a pressing need for localized stimulation to support control of multi-actuator devices with multiple degrees-of-freedom. Integrating the stimulation interface into the device structure may present a robust solution (*34, 49*). Engineering the compliant device structure to compensate for the muscle actuator’s tissue compaction force but minimize the structure’s resistance to active contraction force (e.g., stiffness cancellation (*63*), nonlinear load profile (*61*)) could also allow biohybrid robot designers to better harness the actuator’s force output for the functional goals of their device.

Despite these remaining challenges and opportunities, the novel suspended compliant skeleton approach presented here represents a substantial step forward in the force capabilities and device scale achievable with biohybrid muscle-based actuators. Beyond advancing the capabilities of biohybrid robotics, this platform lays the groundwork for broader applications in tissue engineering, regenerative medicine, and soft robotics. Its modularity and adaptability could facilitate the future development of customizable therapeutic devices, dynamic biological models for scientific research, and next-generation lab-on-chip musculoskeletal or neuromuscular systems. Our results suggest that this approach may serve as a foundational infrastructure for building resilient and high-performance biohybrid machines.

## Conclusion

This work developed a modular approach to culture muscle actuators and construct biohybrid devices using a suspended compliant skeleton for force enhancement during maturation and modular magnetic interfaces for device assembly. The skeleton provides mechanical cues and resistance against the muscle actuators’ spontaneous contraction that enable self-exercise with high stroke lengths. Through their engineered translational degree of freedom, stiffness, and large-stroke design, the compliant skeleton enabled biohybrid actuators fabricated using the accessible, immortalized C2C12 cell line to develop force and stroke capabilities similar to those of actuators fabricated from harvested skeletal muscle cells. Furthermore, the increased force capability broadens the biohybrid robot design space and supports manufacturing mesoscale devices. Once matured, the biohybrid muscle-based actuators can be assembled onto a target device structure through the embedded magnetic interface and released from the compliant skeleton, thereby decoupling the maturation scaffold from the target structure. This modularity further enabled the use of muscle actuators in ensembles to create more complex biohybrid machines. When arranged in parallel, multiple muscle actuators can generate larger force outputs and provide redundancy against partial actuator failure. When arranged in series, the driven device can have an amplified range of motion and multi-DOF mobility. The magnetic interface also enabled replacement of actuators in the event of damage or functional loss. In summary, the presented method enabled the manufacturing of biohybrid devices on a larger scale, with high actuation capabilities, that can be modularly constructed into complex devices, substantially expanding the possible design space in biohybrid robotics.

## Methods Summary

Compliant skeletons were designed with the Freedom and Constraint Topology method (*64*), which kinematically models the relation between the required compliant flexure layout and the desired degree-of-freedom (DOF) mobility. Skeletons were designed to support three actuator lengths (Fig. 2A), allowing for the investigation of the effect of actuator length on stroke and force. By setting translation along the longitudinal axis as the primary DOF, the compliant skeleton was modeled with parallel blade flexures (*65*) perpendicular to the target DOF axis, thereby suppressing other DOF (Fig. 2B). Skeletons were fabricated with UV-curable silicone and 3D stereolithography printing (Fig. 2C). The skeletons can be customized for different stiffness by changing the length, width, and height of the flexures. All actuators were characterized in finite element simulations and through mechanical tests (Fig. 2D, Fig. 2E). Good agreement was found between the experiments and simulations, and the characterized stiffness was used to calculate the actuator force output from the skeleton compression.

We used two skeleton groups to study the effect of skeleton design parameters (length and flexure stiffness) on the resulting performance (Fig. 2F, Fig. 2G). Within both groups, we vary and investigate the impact of actuator length. In the first group, the stiffness of skeletons of varying lengths was not found to vary significantly (*p* > 0.05), yielding an effective stiffness of 2.169 ± 0.146 mN/mm. In the second group, the skeletons have a proportional compliance to their varying lengths. Assuming the muscle actuators have a similar force production regardless of length, the skeletons in the second group will compress proportionally more when the actuators contract, leading to equivalent strain. By comparing actuators of the same length between two groups, we explore the impact of skeleton stiffness on their matured performance.

Our bioink was adapted from the literature (*32, 39*), consisting of C2C12 murine myoblasts at a concentration of 10 million cells/mL, 30% v/v Matrigel, and thrombin and fibrinogen as the cross-linking mechanism. The mixture was injected into a mold with a ring-shaped cutout, whose surface was treated with Pluronic F-127 to prevent cell adhesion, facilitating ring removal and transfer. The injection volume was proportional to the length of the ring, ranging from 145, 185, to 225 *μ*L for the 10, 15, and 20 mm rings, respectively, creating a targeted 4 mm^2^ fiber cross-section. The cast bioink was cultured in supplemented growth media (GM) for three days to allow the C2C12 myoblasts to proliferate and the construct to compact in the mold before being transferred to the compliant skeleton and maturing in differentiation media (DM). Both media were supplemented with 6-aminocaproic acid (ACA) to slow the decomposition of fibrinogen in the matrix (*39*), which can cause the ring to break (*66*) during the early maturation stage before the myotubes were sufficiently developed to provide structural integrity. To balance the yield rate and the remodeling of the hydrogel matrix during actuator maturation, we employed a temporal ACA concentration schedule, which substantially improved manufacturing yield (See Supplemental Text). On the compliant skeleton, the rings were cultured in 3 mg/mL ACA for three days before lowering the concentration through partial media exchanges with DM at sequentially lower ACA concentrations, until reaching 1 mg/mL. This approach led to a 94.898% yield rate on day 14 (D14), the end of the maturation stage for this study. By contrast, the rings cultured in a constant 1 mg/mL ACA concentration started to fail and break on day three and reached a 0% yield rate within seven days in DM. Spontaneous contractions were observed as early as day five for the temporal concentration group, only a day later than the constant 1 mg/mL group. To actuate matured muscle actuators, biphasic electric fields were applied to the surrounding media at 0.353 V/mm, 10 ms pulse width, and oriented along the length of the actuator.

Detailed materials and methods can be found in the Supplemental Materials.

## Supporting information

Compliant skeleton and accessories CAD files

## Acknowledgments

The authors thank Drs. Maria Guix and Rafael Mestre for stimulating conversations about bioink formulation, ACA supplementation, and compliant skeletons. The authors thank Richard Desatnik for sharing the 3D-printed master mold processing technique. The authors acknowledge the use of generative AI during the implementation of the computational analysis tools. The generated scripts were inspected and validated by the authors before use, and the outputs were examined before being included in the manuscript. The prompts used to create the tools are provided in the supplemental materials.

## Author contributions

H.Y. and V.W.W. conceptualized this research. H.Y., A.S.W., J.M.S., D.R., D.M., E.T., and V.W.W. conducted research activities. H.Y., A.S.W., and K.C. curated and analyzed datasets. H.Y., A.S.W., J.M.S., D.R., K.C., and D.K.P. developed the methods. A.W.F., T.C.K., and V.W.W. acquired funding for and supervised this research. H.Y. and V.W.W. were responsible for project administration. H.Y., A.S.W., D.M., and V.W.W. composed the first draft of the manuscript. All authors reviewed and commented on the manuscript.

## Data and materials availability

All data used to create plots and figures are accessible at https://doi.org/10.5281/zenodo.15814979. The scripts for analyzing compliant skeleton compression, spontaneous contraction, and cross-section live/dead images are available at https://github.com/CMU-BORG/compliant-skeleton-muscle-actuator. Detailed methods, data figures, and tables are available in the Supplementary Materials. Computer-aided design models of the compliant skeletons and accessories are available in the supplemental files.

## Supplementary materials

Materials and Methods

Supplementary Text

Figure S1 to S7

Tables S1 to S9

Video S1 to S7

Compliant skeletons and accessories digital models

