## Supplementary materials document for "Modular Assembly of Biohybrid Machines Using Force-Enhanced Skeletal Muscle Actuators"

### Supplementary Materials for
### Modular Assembly of Biohybrid Machines Using Force-Enhanced Skeletal Muscle Actuators

Humphrey Yang[1*], Avery S. Williamson[1], Ji Min Seok[1], Daniel Ranke[3], Duncan McGee[2], Emily Trotto[3], Kiyn Chin[1], Dinesh K. Patel[1] Adam W. Feinberg[2,3], Tzahi Cohen-Karni[2,3], Victoria A. Webster-Wood[1,2,3,4,5*]

[1]Dept. of Mechanical Engineering, Carnegie Mellon University, Pittsburgh & 15213, USA.

[2]Dept. of Biomedical Engineering, Carnegie Mellon University, Pittsburgh & 15213, USA.

[3]Dept. of Materials Science and Engineering, Carnegie Mellon University, Pittsburgh & 15213, USA.

[4]McGowan Institute for Regenerative Medicine, Carnegie Mellon University, Pittsburgh & 15213, USA.

[5]The Robotics Institute, Carnegie Mellon University, Pittsburgh & 15213, USA.

**This PDF file includes:**

Materials and Methods

Supplementary Text

Figure S1 to S7

Tables S1 to S9

Video S1 to S7

Compliant skeletons and accessories digital models

### Materials and Methods

#### Compliant skeleton and hardware fabrication

***Compliant flexure fabrication:*** All versions of compliant skeletons and hanger accessories were designed in Rhinoceros 3D 8 (Robert McNeel & Associates) and SLA printed out of Silicone 40A (Formlabs) on a Form 3+ (Formlabs) using the default manufacturer-recommended settings (0.100 mm layer thickness). Models were sliced in PreForm (Formlabs) with compliant flexures built directly on the platform, aligned along the y-axis in parallel with the laser strafe, and without supports or raft. After printing, parts were removed from the build platform with plastic tweezers and gently wiped with Kimwipes (KimTech) to remove excess resin. Parts were initially washed in a solution of 20% n-butyl acetate and 80% IPA, and sonicated for 15 minutes. This was repeated for a second wash cycle before the parts were removed and laid flat, without stacking, on a petri dish to dry in a chemical hood with an amber light for at least two hours. Once dried, iron-PDMS inserts (for the suspended skeletons, See ***Modular magnetic interface fabrication***) or neodymium magnets (for the compliant devices) were inserted and sealed into the structure by applying a small amount of Silicone 40A to the opening with a fine-tipped syringe (20G). After 5 minutes of rest to allow the ink to settle, it was cured in place by a 10-second flash using a high-power UV torch (UV Beast V3 385-395 nm). Parts were placed in a 6-well plate filled with DI water and cured in a UV flood-curing chamber (Phrozen Cure Station) for 45 minutes. Once cured, the sacrificial supports on the skeleton were removed (Fig. S1A), and the hanger and skeleton were assembled for later use in muscle ring transfer (see ***Assembly of cast rings on suspended compliant skeletons***).

***Modular magnetic interface fabrication:*** To enable magnetic modular assembly of the skeleton backbones on various devices, magnetic inserts were fabricated and assembled into the pillars of the skeleton. To fabricate the inserts, 22 g of iron powder (44890, Sigma-Aldrich) and 5 g of PDMS base (Sylgard 184 A, Dow Corning) were added to a 20 mL glass vial, hand mixed for 2 minutes, and agitated in a planetary centrifugal mixer (AR-100, Thinky) for another three minutes at 2000 rpm. 0.5 g of PDMS crosslinker (Sylgard 184 B, Dow Corning) was added, and the entire solution was mixed again on the planetary centrifugal mixer for three minutes, followed by three minutes of defoaming. The ink was poured into a 35 mm Petri dish, and a 2 mm thick casting mold (Fig. S1B) was lowered and pressed down flat onto the poured ink to fill the 2 mm wide cylindrical holes. The Petri dish was covered with a lid, vacuum desiccated for 15 minutes, then blown with 200 °C hot air with a hot air soldering gun (X-Tronic 5000 Series) for one second to remove small bubbles on the surface. The casted molds were left to cure in a 65 °C oven (Oakton StableTemp Model 05015-51) for 5 hours. To retrieve the iron-PDMS inserts, a scalpel was pushed along the mold surface to remove excess material and make sure the inserts have a 2 mm height. Next, the inserts were pushed out from the casting holes with a plastic tweezer and inspected for imperfections before assembling them onto the compliant skeleton backbones.

***Suspended compliant skeleton culture system accessory fabrication:*** To suspend the compliant skeletons in the wells, thereby eliminating the effect of friction or surface tension on the loading conditions experienced by the actuators during maturation and testing, several custom well inserts and testing frames were created. These additional parts were designed in Rhinoceros 3D version 8 (Robert McNeel & Associates), sliced in PrusaSlicer 2.8 (Prusa Research, Slic3r), and 3D printed out of Polylactic acid (PLA, Polymaker) filament on an Original Prusa MK4 (Prusa Research). The same sterilization technique used for the PDMS casting molds was also applied to preparing the 3D printed components for cell culture.

Muscle ring fabrication

***Stereolithography (SLA) printing of the master mold:*** To fabricate the C2C12 ring casting molds, a master mold was designed in Rhinoceros 3D version 8 (Robert McNeel & Associates) and Stereolithographically (SLA) printed out of Tough 2000 (Formlabs) on a Form 3+ printer (Formlabs) using the default manufacturer-recommended settings. Part models were sliced in PreForm (Formlabs) and oriented 75° from the build plate using minimal supports to promote curing on both sides in the flood UV chamber. After printing, parts were removed from the build platform with supports intact and washed in a FormWash (Formlabs) machine using the default manufacturer-recommended settings. Then, parts were washed twice in fresh Isopropyl alcohol (IPA) and sonicated for 10 minutes each before drying overnight. The next day, molds were flood-cured in a FormCure (Formlabs) using the manufacturer-recommended temperature for 10× the recommended time (600 minutes). Once fully cured, the supports were carefully removed, and the final part was washed in DI water and left to dry thoroughly (Fig. S1C).

***Fabrication of Polydimethylsiloxane muscle ring casting wells:*** Muscle ring casting wells were fabricated using Polydimethylsiloxane (PDMS, (Sylgard 184, Dow Corning)) and the master mold. Prior to casting the PDMS wells, the holes on the back of the master mold were sealed with Scotch tape, and the master mold was placed in a Petri dish. PDMS was mixed at a 10:1 ratio (base:crosslinker) with a planetary mixer (AR-100, Thinky) at 2000 RPM for three minutes of mixing and three minutes of defoaming. The mixture was then poured into the wells of the master mold, ensuring that all geometric features were covered. The casted molds were vacuumed in a dessicator for 15 minutes at room temperature, then blown with 200°C hot air with a hot air soldering gun (X-Tronic 5000 Series) for one second to remove small bubbles on the surface. Next, the Petri dish was covered and left to cure in a 65°C oven (Oakton StableTemp Model 05015-51) for 5 hours before carefully removing the muscle ring casting wells from the master mold.

To prepare the muscle ring casting wells for bioink injection, they were washed in a beaker containing 70% ethanol for three times prior to transferring them to a biosafety cabinet (BSC). For each wash, the container was gently swirled for 30 seconds with the parts completely submerged in 70% ethanol. At the last wash, the well molds were left inside the beaker and remained covered in ethanol while being transferred to the BSC to ensure sterility. There, wells were removed from ethanol and washed individually by dipping and swirling them in a Petri dish containing sterile Phosphate buffered saline (PBS, Gibco) three times (5 seconds each time), then moved to a clean Petri dish. After removing residual PBS from the casting wells using pipettes, the casting cavity was loaded with 1% Pluronic F-127 (Sigma-Aldrich P2443) solution equal to the casting volume to reduce adhesion between the bioink and the casting well in subsequent steps. The casting wells with Pluronic solution in the wells were then left in the BSC for incubation and UV light sterilization for one hour. Finally, the Pluronic solution was aspirated, obtaining a coated and dry well mold for bioink injection (Fig. S1D).

***C2C12 culture and biohybrid actuator ring casting:*** Prior to bioink fabrication, C2C12 murine myoblasts (CRL-1772, ATCC) were cultured in growth media (GM) consisting of high-glucose Dulbecco's modified Eagle's medium (DMEM; Gibco) supplemented with 10% fetal bovine serum

(FBS; Gibco), 1% of 200 nM L-glutamine (Gibco), and 1% of penicillin-streptomycin (10,000 units/mL penicillin, 10,000 $\mu$g/mL streptomycin, Gibco). Cells were cultured in T150 and T500 flasks (Thermo Scientific) at 37°C and 5% $CO_2$, with media changes evey two days. Cells no higher than passage 5 were passaged following the manufacturer's recommended protocols before reaching 50-60% confluency until a sufficient quantity (ranging from 1.45-2.25 million cells per ring, depending on the ring length) of cells was reached for casting (Fig. S1E).

To create the biohybrid muscle-based actuators, a casting bioink was mixed from two precursor solutions. The bioink formulation was adapted from Raman *et al.* (*15*) and Guix *et al.* (*39*) with slight modifications. C2C12 cells, having reached the target cell quantity and 80% confluency in T-500 flasks, were collected by trypsinization. Briefly, GM was aspirated and the flasks were washed with 1× PBS (Gibco). The PBS was aspirated from the flasks, and the PBS wash was repeated twice. After removing the remaining PBS from washing, 0.25% Trypsin-EDTA (Gibco) was added to each flask and then the flasks were incubated at 37°C and 5% $CO_2$ for 5 minutes. GM was then added to the flasks to stop trypsinization. Supplemented growth media (GM+) was made by supplementing GM with 3 mg/mL of aminocaproic acid (ACA, Sigma-Aldrich). The fibrinogen stock solution (120 mg/mL) was made by dissolving fibrinogen (Sigma-Aldrich) in PBS. Following centrifugation and cell counting, the cell pellet was resuspended at a concentration of 20 million cells/mL in an 8 mg/mL fibrinogen solution and then vortexed to form Part A of the bioink. Lyophilized thrombin powder (Sigma-Alrich) was dissolved in 0.1% (w/v) bovine serum albumin (BSA, Thermo Scientific). Part B of the bioink consisted of 30% (v/v) Matrigel (Corning) and 4 units/mL thrombin in GM+. Part B was made by first mixing the thrombin and GM+ and vortexing, and then Matrigel was carefully pipetted into the mixture. Part B was carefully mixed by aspirating up and down while minimizing the introduction of bubbles. To form the complete bioink for casting, equal volumes of Part A and B were quickly but carefully mixed by pipetting up and down, again trying to limit bubbles. The resulting cell-laden bioink had the following final concentrations: 10 million cells/mL, 4 mg/mL fibrinogen, 30% (v/v) Matrigel, and 2 units/mL thrombin in GM+. The cell-laden bioink was quickly pipetted into the PDMS casting wells located in a 6-well plate (VWR Caplugs, Fig. S1F). The rings were incubated for 30 minutes at 37°C and 5% $CO_2$ to allow the Matrigel and fibrinogen mixture to gel. GM+ was then added to each well of the 6-well plate to cover each casted ring (Fig. S1G). Partial media changes were performed daily for two days (on D-2 and D-1) until ring transfer (D0) to the compliant skeletons.

Biohybrid muscle-based actuator maturation on suspended compliant skeletons

***Assembly of cast rings on suspended compliant skeletons:*** After compaction, the C2C12 rings were transferred to the compliant skeletons for maturation in differentiation media (DM+) consisting of high-glucose Dulbecco's modified Eagle's medium (DMEM; Gibco) supplemented with 10% horse serum (Gibco), 1% of 200 nM L-glutamine (Gibco), 1% of penicillin-streptomycin (10,000 units/mL penicillin, 10,000 $\mu$g/mL streptomycin, Gibco), and 0.05 $\mu$g/mL of insulin-like growth factor 1 (IGF-1; Sigma-Aldrich) supplemented with varying concentrations of ACA (Sigma-Aldrich) starting at 3 mg/mL and tapering to 1 mg/mL (See Table S4).

In the six-well plate containing the muscle ring construct and casting molds, the molds were gently lifted from the bottom of the well with the muscle ring construct inside and flipped upside-down so that gravity could aid in the removal of the rings from the molds. The rings were gently displaced with forceps, first at the interface with the outer wall of the mold, and then at the interface with the inner pillars, such that the ring was carefully worked off the pillars evenly around the circumference. In particular, the muscle rings were displaced off the pillar at equal pace across its length to minimize stretching and twisting. Once fully removed and free-floating in the culture well, a tweezer was inserted through the ring, and a second pair of tweezers was used to gently nudge the ring onto the first pair. To ensure safe transfer from the well plate containing the casting model to the well plate containing the compliant skeleton, the second pair of tweezers should hold onto the first pair of tweezers' tip, such that should the muscle ring slide along the first pair of tweezers, the muscle ring will rest onto the second pair of tweezers instead of falling off. It should be noted that any pinching of the ring should be strictly avoided as this can induce weak points that risk actuator failure during maturation.

In the maturation well plate, each well is prepared with a single skeleton by adding 7 mL of DM+ containing 3 mg/mL ACA. Prior to muscle ring mounting, the skeletons should be prepared and placed in a Petri dish with the pillars facing down and the hanger accessory attached on top of the skeleton. Once the muscle ring constructs were transferred to the maturation well plate, they were left lying at the bottom of the wells. The skeleton was then lifted using a pair of tweezers and gently compressed in the longitudinal direction and lowered toward the muscle ring inside of the well. Once the pillars on the compliant skeleton were inserted through the opening of the ring, the skeleton should be gently released to allow the notch on the pillars to hold onto the muscle ring fiber without introducing damage. The skeletons, with the muscle ring on top, were then gently agitated to ensure the rings were secured in place. The 3D printed inserts were then assembled onto the well plate, and the entire skeleton was mounted onto the insert and suspended from the top of the well to elimnate friction between the skeleton and bottom of the well (Fig. 1A). Once mounted in place, the hanger accessory's sacrificial frame was cut away using a pair of stainless steel scissors (Fig. S1H). The suspended compliant skeleton actuator assemblies were subsequently cultured at 37°C and 5% $CO_2$.

***Periodic imaging and media changes during maturation:*** During tissue maturation, the assembled actuators (Fig. S1I) were imaged daily in their wells using an Echo Revolution Microscope (Discover ECHO - Bio Convergence Company). The microscope was configured to the inverted mode, such that the camera could inspect the actuator from the bottom without the hanger accessories obscuring the view. The microscope's multi-point imaging function was used to create a stitched image for each well. The focal distance was adjusted to be in plane with the muscle rings, with the movement speed set to 3% and the acceleration level to slow to minimize shaking the skeletons.

After imaging, the well plate was sprayed down with a 70% ethanol solution and transferred back to the BSC. Inside the BSC, the lids were carefully removed to aspirate 3-4 mL of media from each well, depending on the ACA tapering schedule (Fig. S1K). 3-4 mL of fresh, warmed DM+ with the appropriate ACA concentration (See Supplemental Text) was added to each well before closing the lids and replacing the well plate in the incubator at 37°C and 5% $CO_2$. For the first three days of culture, partial media changes were performed by removing 3 mL of media and replacing it with 3 mL of DM+ supplemented with 3 mg/mL ACA to maintain the high initial ACA concentration. Starting on Day 4, media was replaced with 3 mL of DM+ with decreasing concentration of ACA (Table S4). Beginning Day 8, due to increasing media consumption and pH decreases between media changes, the amount of the partial media change was increased to 4 mL of DM+ supplemented with 1 mg/mL ACA. This rate of media change and concentration was maintained for the remainder of the culture until actuators were tested at Day 14. To assess the effect of the time-dependent ACA concentration taper on manufacturing yield, rings were fabricated using the same methods described above and cultured in media with a fixed concentration of 1 mg/mL ACA for all 14 days (Fig. S1K). Yield rate for both treatments was calculated based on the number of actuators intact each day at the time of media change.

Biohybrid muscle-based actuator stimulation and force characterization

Prior to testing, the biohybrid actuators were imaged on D14 using the same protocol as described above (Fig. S2). A pair of tweezers and the stimulation hardware were sterilized in 70% ethanol solution for a minimum of five minutes and allowed to dry in open air. DM+ with 1 mg/mL ACA was prepared in a volume corresponding to the number of actuators to be tested (7 mL/actuator), aliquoted into 50mL conical tubes, and heated in a 37°C water bath. Immediately before testing an actuator, a single well in a fresh 6-well plate was flooded with the warmed DM+ 1mg/mL ACA solution, and the conical stock media tube was returned to the water bath to maintain temperature. Actuators were tested one at a time with the following procedure. An assembled well insert carrying a ring-laden compliant skeleton was inserted into the open well, covered with a lid, and transferred to the microscope for continuous imaging with a 1.25× lens during stimulation. Once secured on the microscope such that the edges of the skeleton were aligned with the y-axis of the viewport, the lid was removed, and the stimulation insert was installed such that the mesh electrodes (platinum-coated titanium mesh electrode, Redoxme AB) on the stimulation attachment were aligned with the ring's longitudinal axis (Fig S3). The mesh electrodes had a width of 5 mm and thickness of 0.5 mm. The electrode was cut into an appropriate length such that when assembled, the tip of the wire mesh was submerged and touched the bottom of the well. A 26 AWG wire (Calmont Wire & Cable Inc) connected to the opposite end of the mesh electrode by exposing and tightly wrapping the metallic core around the mesh. The distance between the electrodes was measured to be 28.32 mm.

Once video capture was configured and initiated, the actuator was stimulated using a High-Power, Bi-Phase stimulator (Aurora Scientific, 701C). The pulse phase was configured to Bi-phase, the range set to 20V, and adjusted to 50% to provide 10V stimulation across the electrodes, creating an electric field of 0.353 V/mm. Stimulation trains were controlled using Aurora Scientific's Dynamic Muscle Control LabBook software using the Instant Stim mode with a pulse width of 10 ms. In the configurations, the pulse count was set to ≥ 999 to saturate the 6-second stimulation window, and the training frequency, which allows the instrument to create repeated stimulation windows, was set to 0.001 Hz (i.e., once every 1,000 seconds) to avoid repeated stimulation during the test period. Each 6-second stimulation period was followed by a 24-second rest period for a total stimulation cycle of 30 seconds. Table S1 details a typical stimulation schedule with multiple cycles from 1 Hz to 50 Hz. See Fig. S4 for representative images of stimulated skeleton compression.

After the stimulation test had concluded on a given actuator, the next well in the 6-well plate was prepared with fresh, warmed DM+, and the protocol was repeated with a new actuator. Once all scheduled tests were complete, the rings were transferred to a new 6-well plate filled with 7 mL of PBS and gently swirled to wash. PBS was aspirated from the plate with a 10 mL pipette, replaced, and the wash process was repeated. The rings were then fixed for IHC imaging with the addition of 7 mL of 4% paraformaldehyde-PBS solution (ThermoFisher Scientific) and sealed with Parafilm (ULINE) to store in a refrigerator at 4°C.

Devices assembly, repair, and characterization method

The devices shown in Figure 5 and Figure 6 were fabricated with the same method as the compliant skeletons, and they were cleaned by submersion in 70% ethanol for five minutes and let dry in air before use. The experiments were conducted in 100 × 15 mm square Petri dishes (Electron Microscopy Sciences) filled with Phenol red-free DM+. The differentiation media was prepared with the same composition as that used for muscle ring maturation, except for using phenol red-free DMEM (31053028, Gibco) instead of regular DMEM. To assemble the devices, matured muscle actuators and compliant skeletons were freed from the hangers and assembled onto the devices via the magnetic interface using plastic tweezers. A pair of ceramic scissors (Slice) were then used to cut the neck connecting the pillars and the compliant skeleton, releasing the muscle actuator from the passive resistance provided by the compliant skeleton. The stimulation method was identical to that used for muscle actuator characterization. To stimulate the muscle actuators, a pair of mesh electrodes (platinum-coated titanium mesh electrode, Redoxme AB) of appropriate size (i.e., spanning the width of the actuators) were placed in alignment with the muscle actuators where possible. For the 2-DOF positioning stage, the electrodes were placed along the x-axis since it was not possible to align with both actuators simultaneously due to their orthogonal arrangement. The stimulation voltage varied between setups and was calculated based on the distance between the electrodes to create an electric field of 0.353 V/mm. All devices were able to recover to their rest pose without manual reset with the exception of the serial robotic arm-like device, where the static friction between the end effector and the bottom of the Petri dish prevented the compliant skeleton from springing back. To reset the device, we manually pushed the end effector and allowed it to settle with the spring force of the joints.

#### Live/Dead and Immunohistochemistry Imaging Sample Preparation

***Immunohistochemistry:*** Immunohistochemistry imaging was performed on muscle rings targeting myosin heavy-chain, actin, and nuclei. Directly before antibody staining, the muscle rings fixed at the end of force characterization were carefully removed from their skeleton, immersed in 0.2% Triton X (Sigma, X100) for permeabilization for 30 minutes at room temperature, and treated with an albumin blocking solution (1% w/v in PBS, Sigma, A1653) for 1 hour. After washing, rings were fully immersed in a 1:200 solution of Myosin Skeletal Muscle Monoclonal Antibody (in PBS, Thermo-Fisher, MA5-11748) and gently agitated at 4°C for 48 hours. Following a triplicate wash in cold DPBS, rings were transferred to a solution of 1:200 CoraLite Plus 488-conjugated Alpha Actinin Polyclonal antibody (Thermo-Fisher, CL488-11313), 10 $\mu$g/mL Hoescht (62249, ThermoFisher Scientific), and 1:500 Donkey anti-Mouse IgG (H+L) Highly Cross-Adsorbed Secondary Antibody to label the myosin skeletal muscle primary antibody (Thermo-Fisher, A-31571) for an additional two days at 4°C with agitation. The muscle rings were then washed three times in DPBS and stored at 4°C in DPBS until imaging.

All muscle ring immunohistochemistry imaging was performed using a Nikon AXR confocal microscope with either a 10x air (PLAN APO OFN25, NA=0.45) or 60x oil immersion objective (PLAN APO VC 60x, NA=1.20). Images were all adjusted with NIS-Elements AI-assisted noise correction and displayed as maximal intensity projections of Z-stack images of mid-ring sections in both low and high magnification conditions.

***Cross-section live/dead imaging:*** On D0 and D14, muscle ring samples (n=3) for live/dead imaging were removed from their skeletons, washed three times in 37°C Dulbecco's phosphate buffered saline (DPBS, Gibco), followed by staining with a mixture of calcein and ethidium homodimer-1 (Invitrogen, L3324) with Hoescht (10 $\mu$g/mL, 62249, ThermoFisher Scientific) at 37°C under low agitation for 90 minutes. After the staining process had concluded, rings were washed again three times in 37°C DPBS, followed by immobilization in 1.5% w/v saline-formulated agarose (Invitrogen, 16500-100) and 37°C. Once the agarose solidified, the muscle rings were transferred to a bath of ice-cold PBS for slicing with a Leica VT100S vibratome. Each muscle ring was cut into 300 $\mu$m slices at 90 Hz frequency and approximately 50 $\mu$m/s cutting speed, with 6-12 slices generated per ring. These slices were immediately imaged on a Nikon AXR confocal microscope under a 10x objective (PLAN APO OFN25, NA=0.45) for quantitative live/dead studies. Whole slices were reconstructed through a stitched image with 25% overlap for proceeding analyses.

***Cross-section live/dead analysis:*** A custom Python script was used to analyze the cell distribution across the cross-section. The cross-section was divided into several 100 $\mu$m bins based on their depth from the cross-section perimeter. Each depth bin was characterized by its mean signal intensity per area $I = \frac{\sum V}{A}$, where $\sum V$ sums the pixel values of a channel (i.e., nuclei, live, or dead) in a depth bin, and A is the total area of the depth bin (in $\mu m^2$). By using the outermost bin of depth 0-100 $\mu$m as the baseline, we could then calculate the change in signal intensity with $I_{relative} = \frac{I}{I_{0-100}}$. A $I_{relative}$ lower than 1 indicates less signal than the outermost bin of the cross-section, and vice versa. Since each cross-section image may have a different number of cells and signal strength (pixel value), directly pooling the raw measurements may lead to large deviations. By contrast, the relative signal approach normalized each image against its own baseline to produce a unit-less value that represents the signal change versus depth, thereby allowing more robust data-pooling and statistical analysis across multiple images.

Compliant skeleton stiffness characterization

***Compression testing assembly:*** Compliant skeleton stiffness was analyzed on a universal testing system (MTS Systems, Criterion Model C41) with a biobath accessory using a 5N load cell with a +/- 0.05% applied force precision error. The test samples were fabricated with the same process as the compliant skeletons used for cell culture, but they were modified with end tabs for mounting on the test instrument. To ensure that the samples were mounted in an unstrained loading condition, they were assembled into a custom 3D-printed (Prusa MK4) PLA frame that held them at rest length using sacrificial spacers until they were ready to be tested. For each sample, the end tabs of the skeleton were initially secured to the spacer side of the frame with double sided tape (Fig. S5A). The matching ends of the frame were then mounted over the skeleton and bolted together with the spacer (Fig. S5B). Silicone sacrificial support bars were then removed from the skeleton flexures with a scalpel (Fig. S5C) to allow for free movement of the flexures. The lower end of the frame was slotted onto the lower mounting bracket in the MTS biobath and bolted in place (Fig. S5D). The MTS crosshead was lowered slowly into slotted contact with the upper frame end until bolts could be inserted through all components to fix the frame's position without exerting any load on the crosshead (Fig. S5E). This position was recorded as the zero extension position. Then, the PLA spacer beams were cut and removed to uncouple each side of the frame and allow for deflection of the skeleton (Fig. S5F). The biobath was sealed and filled with DI water at room temperature to a fixed height before recording the zero load condition (Fig. S5G).

***Cyclic compression protocol:*** Once assembled into the testing assembly, TestSuite Elite (MTS Systems) was used to detect the measured load and extension of the skeleton. The 10, 15, and 20 mm skeleton samples (N=3 each) were compressed by 2.5 mm, 3.75 mm, and 5 mm, respectively, at a 10 mm/min strain rate for 10 cycles with a 10 Hz sampling rate. Tests were repeated at each length with the mounting hardware and without samples to measure the ambient buoyancy forces. Load vs Extension data was recorded to a CSV and subsequently analyzed in MATLAB.

In MATLAB R2024b (MathWorks), the load vs extension data was initially resampled at 10 Hz and lowpass-filtered with a cutoff frequency of 1 Hz, steepness of 0.99, and a stopband attenuation of 60 dB. Each dataset was fit to the linear equation $y = a * x + b$, where the slope a was recorded as an equivalent stiffness. A given sample was filtered, fitted, and then corrected for buoyancy before calculating the effective stiffness. Figure S5H shows an example of this analysis for determining buoyancy forces. The mean stiffness and standard deviations were calculated for each A-F sample group (Table S2). The $R^2$ values for all samples based on linear fitting are also reported for both pre- and post-filtering (Table S3).

#### Supplementary Text

Tapering ACA concentration during maturation improves biohybrid actuator yield

To investigate the effect of tapering ACA concentration on manufacturing yield, we compared the yield rate of rings cultured with a fixed 1 mg/mL ACA concentration ($N = 23$ actuators) as has previously been reported in the literature (*39, 66*) to that of the actuators cultured following our ACA taper schedule, where the concentration was asymptotically decreased from 3 mg/mL at differentiation onset to approximately 1 mg/mL by Day 14 when the actuators were tested (Table S5). Yield rate was calculated for all batches of actuators used to generate the main findings in this work, resulting in a total of $N = 98$ actuators cultured with the taper. In the group cultured with constant 1 mg/mL ACA supplementation, actuators exhibited high failure rates. For the first 3 days of differentiation, all actuators remained intact, but by the third day, 16 out of 23 actuators had snapped. Of the remaining actuators, all were broken by day 7 of differentiation, resulting in a final yield rate of 0%. In contrast, actuators matured with the ACA taper schedule demonstrated high yield. In all experiments, all actuators remained intact until day 7. Few actuators broke between day 8 and 14, resulting in a final overall yield rate of 94.898% when using the ACA taper.

Muscle actuator fiber cross-section live/dead analysis.

Our cross-sectional analysis revealed that the muscle actuator had a necrotic and cell-void core, as previously observed in similar biohybrid muscle-based actuators (Fig. S6). The live cells were primarily located around the perimeter of the fiber's cross-section under the 200 $\mu$m oxygen diffusion limit (Fig. S7), and the number of live cells decreased with depth. Noticeably, the core of the muscle actuator was devoid of cells. We speculate that this is due to the migration or death of the centermost cells from hypoxia following bioink injection. Comparing D0 and D14 images, the cross-sections were also qualitatively observed to become more circular throughout the maturation period, and the relative distribution of cells across the cross-section also became more disparate.

Using finite element method to simulate compliant skeleton mechanical properties.

***Simulation setup and parameters:*** All finite element simulations were completed in Ansys 2024 R2 using static structural simulation. The silicone 40A constituting the compliant skeletons and devices was modeled as an isotropic material with a specific weight of 1.02 $g/cm^3$, elastic modulus of 1.02 MPa, and a Poisson's ratio of 0.495 by assuming a rubber-like incompressibility. The simulation input geometries were modeled as an assembly of compliant flexures and rigid bodies connected by rigid contact interfaces. To balance computation time and mesh resolution, the compliant flexures were discretized with a 200 $\mu$m element size to accommodate the large defelctions commonly observed in flexural systems, whereas the rigid bodies had a 500 $\mu$m element size to reduce computational workload.

To model the compression of a compliant skeleton under the forces exerted by the muscle actuator, one side of the pillar was set as the fixed end, while the other was subjected to a displacement or force load boundary condition. The simulated load curves were acquired by applying an increasing load, plotting the force and displacement measurements, and linearly interpolating between data points.

***Elephant's foot calibration:*** The stereolithography printer used overexposure to ensure adhesion between the printed part and the build plate. This feature also caused the print's bottom layers (up to the 0.7 mm height) to become wider than the intended dimensions and is commonly referred to as the elephant's foot problem. To model this effect, we calibrated our simulation models by modeling the widening. A simulation of the 10 mm compliant skeleton without any modification to the geometry was used to establish a baseline for the stimulated stiffness. The baseline is then compared against the measurements acquired through physical stimulation, which was much stiffer due to the presence of the elephant's foot. The stiffness difference could be interpreted as the ratio between the two cross-sections' second moment of area: $K_{diff} = I_{exp}/I_{baseline}$, where $K_{diff} = K_{exp}/K_{baseline}$ is the difference between the stiffness of the physical and baseline compliant skeletons, $K_{exp}$ and $K_{baseline}$, respectively. Similarly, $I_{exp}$ and $I_{baseline}$ are the respective second moments of inertia of the experimental and baseline flexure cross-sections. Given it was obtained through physical experiments, $I_{exp}$ was modeled with elephant's foot widening as $I_{exp} = \frac{h_{upper}b^3}{12} + \frac{h_{lower}(b+w)^3}{12}$. Specifically, the cross-section was divided into two rectangular components: the upper part $h_{upper}\times b$ was absent of the elephant's foot and the lower part $h_{lower} \times (b + w)$ was widened by overexposure for a width of $w$, thus each edge was offset by a distance of $o = \frac{w}{2}$. The summation $h_{upper} + h_{lower} = h$ should equate to the height of the flexure $h$. Conversely, $I_{baseline}$ was directly calculated as

$I_{baseline} = \frac{hb^3}{12}$.

To compute $I_{exp}$ and $I_{baseline}$, $h = 0.8$ and $b = 0.35$ could be acquired from the CAD model, and $h_{lower} = 0.7$ was obtained from the slicer setting, making $h_{upper} = 0.8$. The elephant's foot widening $w$ could then be solved by plugging $I_{exp}$ and $I_{baseline}$ into $K_{diff} = I_{exp}/I_{baseline}$, making $w = 0.192$ mm and $o = 0.096$ mm. The parameters and results are documented in Table S6.

Computational analysis of muscle actuator performance and maturation

***Skeleton compression tracking:*** A computer vision-based tracking tool was implemented to detect compliant skeleton compression in recorded videos. The tracking tool was implemented in Python version 3.12.2 and leveraged OpenCV for the bulk part of its processing pipeline. Given an input video in either .mov or .mp4 format, the user should tag feature points in the video to initialize the tool. The script also provides a function to calibrate the pixel to $\mu$m parameter based on a known measurement in the video. The script produced a video overlaid with the tracked features and a .csv file recording the distance and position of the tracked features.

The tracking mechanism was based on the Lucas-kanade Optical Flow algorithm provided by OpenCV. The tracking feature was specified by a square window of user-specified pixel dimensions. When using the script, the tool facilitated this with an interactive viewport, allowing the user to pick and place the tracking windows and specify the skeleton inner edge positions when tracking skeleton compression. The script also included a feature recovery function using OpenCV's *matchTemplate* function that re-calculates the tracking window's position should the optical flow algorithm lose track of the features' positions.

To use the tracking tool for detecting skeleton compression, the user should specify two feature windows per side of the skeleton. The displacement of each side of the skeleton is then calculated based on the averaged displacement of the corresponding features. In addition to the features, the user may also specify an edge aligned to the inner edge of the skeleton, whose positioned is updated by applying a static offset from the tracked features. The skeleton compression is subsequently calculated by subtracting the measured edge distance from that measured on an uncompressed compliant skeleton.

***Spontaneous contraction data analysis:*** Spontaneous contraction data were calculated from skeleton compression readouts acquired by the tracking tool. To filter out videos where the muscle actuators were not exhibiting spontaneous contraction, we ignored videos whose range of skeleton compression was smaller than ten times the pixel length (i.e., 44.656 $\mu$m). For the remaining videos, their tracked time-series skeleton compression data were processed with Fast Fourier Transformation to decompose their signal in the spectral domain. The effective range of contraction frequency was clipped by the length of the video and the framerate. The upper frequency limit was set to half of the video framerate (i.e., 15 Hz), whereas the lower limit was set by $Hz = 2/(video\_length)$. When reporting the spontaneous contraction data, we assumed that the muscle actuators may exhibit different modes of spontaneous contraction (i.e., at different stroke length and frequency) simultaneously, and the observed compression was the result of the composition of different spontaneous contraction modes. Thus, the three most prominent spontaneous contraction frequency data was reported in Figure 3A-C. The maximum spontaneous contraction stroke (Fig. 3D) was calculated as the absolute difference between the maximum and minimum compression observed in the video.

***Live/dead cross-section analysis:*** A tool was designed to allow for repeatable analysis of live/dead (L/D) cell distribution across cellularized structure cross-section images. The input images were 3-channel color images of stained cells captured via microscopy, where red corresponds to dead cells, green to living cells, and blue to nuclei. First, the tool finds a contour around the perimeter of the imaged structure, then counts the spatially averaged intensity of red, green, and blue light as a function of distance from that perimeter (i.e., depth into the cross-section).

The tool operates in two modes, single file or batch by folder. When started, the user will be prompted to select between these two modes by a command-line dialog. A GUI interface will then open for selecting either the target image file or a folder containing the batch of target image files, depending on the mode. Then the analysis will run for all selected files. Analysis has an initial interactive phase where the user will use a GUI to tune parameters for each image to find a contour, then a processing phase where all images will be analyzed without user input.

To calculate the contour, first, the image was converted to a binary image using OpenCV's *adaptiveThreshold* method. The binary image was then put through a morphological opening operation. The set of contours was then found via OpenCV's *findContours* method. The contour with the largest area from this set was chosen as the contour around the perimeter of the structure. The user was shown this contour iteratively and can adjust it by the OpenCV trackbars displayed on the UI. The trackbars control the *block_size* and $C$ variables of the *adaptiveThreshold* method, the *kernel_size* used in the structuring element for morphological opening, and the *iterations* of the morphological opening operation. Once the contour was satisfactory, the user could move on to the next image by pressing the "n" key. Once all images had satisfactory contours (Fig. S6), the parameters were noted and stored. Contours were optionally spatially smoothed via a 1-dimensional Gaussian blur in coordinate space.

To count cell signals across a cross-section given a contour, first, a mask was created to indicate all pixels that are inside the contour for subsequent operations. A distance map (i.e., as a single-channel image) was created and initialized to 0, which indicated the distance from the contour of all pixels in that image. Distances were found via a floodfill algorithm with 4-connectivity, starting with the pixels at the contour having a distance of 0, and each successive neighbor increasing that distance. A breadth-first search algorithm was used to identify successors for the flood-fill. Due to banding artifacts in the raw distance map produced via this graph-based method, and the knowledge that adjacent pixels should have correlated distances, the distance map was optionally smoothed via a Gaussian blur. Once this distance map was created, the mean intensity across red, green, and blue channels for a given depth bin was calculated, and L/D graphs as a function of depth can be created. Specifically, the mean intensity $I = \frac{\sum V}{A}$ calculates the average pixel signal in the depth bin, where $V$ is the pixel value of each pixel in the depth bin, and $A$ is the area of the depth bin. To normalize measurements, the outermost bin of depth 0-100 $\mu$m was used as the baseline, and the other bins' signal was normalized against it using $I_{relative} = \frac{I}{I_{0-100}}$. Using this equation, the outermost layer will always have a relative intensity of 1, and decreases toward the center should the core of the muscle fiber have fewer cells.

There were global parameters that affect all images in a given processing run that were set within the code, these are *scale*, *smoothing*, *bin_width*, and *px_to_um_base*. The value of *scale* is a float between 0 and 1 that downscales the images by that fraction in order to speed up processing time, which varies with the square of the scale. While results would technically vary across scales, observed differences for tested images were small. The value of *smoothing* determines the smoothing of the contour as well as the distance map itself. A value of 0 corresponds to no smoothing. The *bin_width* variable controls the contour distances that are binned together, and *px_to_um_base* controls the conversion from pixels to distances at scale 1. The output of the processing will be saved to a dynamically created subfolder of the directory of the Python script. The outputs for an input image ({base_name}.png) will be the parameters selected by the user during the interactive phase ({base_name}_params.json), an image of the resulting contour overlaid on the input image ({base_name}_contour.png), an image of the pixels that are processed based on the contour ({base_name}_mask.png), an image of the distance map ({base_name}_distance_map.png), a line graph of the dead, live and nuclei mean intensity as a function of distance from the external contour ({base_name}_line_plot.png), a bar graph of the same data binned by ranges of distances and converted to fold-count instead of raw intensities via normalization by the counts in the most external bin ({base_name}_fold_change.png), a tabular form of the binned and normalized data ({base_name}_binned.csv), and a tabular form of the original data ({base_name}_full_data.csv).

The cross-section images used to generate the plots in Figure 3I are provided in Figure S6, and the parameters that were used in generating the contours are provided in Table S7. To analyze these images, the *scale* was set to 0.25, *smoothing* to 1, *bin_width* to 100 $\mu$m, and *px_to_um_base* to 0.575.

Generative AI prompts for implementing the analysis pipeline.

***Note:*** The scripts were composed with the help of generative AI in an iterative development strategy. Below are the prompts used (omitting error lookups and format request prompts) . Non-interactive, offline development phases are identified, at which point large language model was prompted by upload of a new version of script.

***Spontaneous contraction analysis:*** Prompts (model version: ChatGPT 4.1):

1. Generate a Python script based on this description. Use camel case and functionalize the script. Global parameters should appear at the top of the script. I have a folder containing multiple .csv files of time series measurements. The time is logged in the ”Second” column, and the measurement is stored in the ”Line distance (um)” column. The measurement is a beating signal. I want to find the frequency and amplitude of the beating signals. When loading each file, check the range of signals first. If the range is smaller than a threshold,

skip that csv file. Next, process the CSV using two methods for comparison. The first method is to use the FFT to find dominant peaks and amplitude. The range of frequencies to report should be based on the length of the video and the frame rate. The upper limit is 30, and the lower limit should have a period of twice the max value in the Second column. Find and report the three most prominent frequencies and amplitudes. The second method is to count peaks in the video. The frequency is then defined by the number of peaks divided by the length of the video, and the amplitude is just the peak's rise. Store the results in a new CSV file that contains the results from all qualifying CSV files in the folder. The output CSV should contain file names, the range of signals, the time length of the data, the number of count peaks, and an entry for each dominant frequency and amplitude. Also, plot the FFT and peak-counting result in the same plot (top and bottom plots) for each CSV file. The FFT should highlight the three peaks, and the peak counting should plot the time sequence with the peaks highlighted. The plots should be saved to a target folder.

2. The measurements are inverted, so the peaks appear as valleys.

3. My input CSV files have a formatted name. P{d}_S{d}_D{d}. Can you order the result CSV so that it's first sorted by D, then by P (i.e., the higher D-value entries will be at the end of the file).

4. Offline Phase: changing FFT frequency clip range and stroke calculation method, formatting, and clean up.

***Muscle actuator cross-section live/dead analysis:*** Prompts (model version: Deepseek R1):

1. Offline Phase: initial script writing, upload 1

2. Modify this python file to have an interactable UI. There are two phases where UI is needed, adjusting the parameters for finding the main contour of the cluster of cells, and adjusting the parameters for finding the contours of each individual cell. These parameters are the threshold, the parameters for the open operation, the kernel size. Create sliders that will on the fly update these parameters and display the resulting contours so the user can adjust till things look right.

3. Offline Phase: feature addition, refactoring and rewriting, upload 2
4. Starting from this python script, update to label pixels by their Manhattan distance from the rasterized contour called depth_map. This is all 0s except for the pixels on the contour. Pixels on the contour have color 1. Do not directly calculate distance, instead Use a variant on the flood fill algorithm that iterates and uses 4-way connectivity recursively until all pixels have nonzero value. On each call, 0 pixels next to nonzero pixels have their value set to (that pixels value + 1)
5. Calculate the max distance between the contour and the centroid, and do not add points to the queue if their distance would be larger than that
6. Use the mask method, rather than the max dist. Create a separate mask that is just the interior of the contour.
7. Switch to adaptive thresholding for finding the main contour
8. There are artifacts in the distance due to noisy contour. How can I smooth the contour before doing flood fill to prevent this?
9. lets use numpy 1d gaussian blur
10. Offline phase: feature addition, refactoring and rewriting, upload 3
11. Starting from this version add the following features. 1. Bin the distances in the output into ranges of 50 um, and use seaborn to display a bar graph, in addtion to a line graph with the un-binned values. 2. Add a toggle to switch between the adaptive thresholding and flat thresholding methods.
12. normalize binned data by the value of the first bin, to produce n-fold measurements
13. save the pandas dataframe to file based on the input fname
14. add the parameters of the trackbars for replicability
15. Offline phase: feature addition, refactoring and rewriting, upload 4

16. Starting from this version, add the following features.
17. Separate the user input phase and the subsequent processing into two phases
18. Upon startup, the script asks the user to select a PNG file or a folder. Create a list of file names, either the PNG file or all the PNG files in the selected fold. For all these files run the user input phase first to select parameters. Each parameter should be stored in a dictionary that indicates which file it is for. After all parameters are established, the run all of the processing.
19. Create an output folder that has a name that encodes the scale and smoothing parameters used for that run. In that output folder, save the csv files, the json file, the raw line graph, the fold_count graph, the input image with the final contour, the smoothed mask, and the visualized depth map. in short, all the outputs of processing. make sure file names encode the name of the input file.
20. rather than labeling the bar graph with bin centers, I want to label with a string that is bin_start - bin_end
21. the rgb_full_data csv is is missing data. When distance is 0, distance bin has no value
22. that didn't work
23. Offline phase: refactoring and rewriting

***Compliant skeleton compression tracking:*** Prompts (model version: Deepseek R1):

1. Offline Phase: initial script writing, upload 1
2. Starting from this python script, write a script that uses roi tracking to determine the distance between two vertical bars over time. When the script is run, the first frame of the video should be displayed. 4 squares and two vertical lines should be drawn, these elements should be able to be clicked and dragged. The user will drag the squares over the corners to be tracked, and the vertical lines will be placed in the middle between the two pairs of boxes. The squares will be used as the rois, and the vertical lines will simply be drawn to maintain the relative horizontal distances between the lines and the average x positions of the two pairs of boxes.

The csv should show the horizontal distance between the two pairs of boxes and the two vertical lines, at every frame of the video.

3. Instructurions for what to press should be in the window title. Also, a version of the video with the tracked objects should be saved in the same place as the csv with similar name. Also the squares should change size based on scroll wheel.

4. The pairs of squares should have different colors so that the user knows which ones to put on the left and which ones to put on the right

5. Instead of the scrollwheel for each individual box, sizes for all boxes should change together, based on the w and s keys. The user should indicate completion with the d key

6. If tracking is lost, save the video anyway. Also color the lines to be matched to their closest box pair.

7. Offline Phase: feature addition, refactoring and rewriting, upload 2

8. This is the current code. When tracking is lost, how can the original template be used to reacquire tracking? Are their methods designed for tracking through occlusion that can be applied?

9. add the core ideas to the most recently uploaded file.

10. use the relative location to the partner in the pair of squares and the last valid position as a guess for tracking recovery.

11. Template matching should be the primary recovery method, look for the original template on every frame. In fact, instantiate a secondary set of squares that just tracks without updating the template, and track both the static template and updated template versions.

12. Offline Phase: feature addition, refactoring and rewriting, upload 3

13. This is the current version that works the best. Try to edit this so that the vertical distance between pairs stays mostly the same, it should be tracked but not very aggressively, i.e. the model in a kalman filter assumes it is constant

14. Offline Phase: feature addition, refactoring and rewriting

Muscle actuator performance benchmark

The plot shown in Figure 1C presented a comparison between unitary muscle actuators that appeared in the literature. Since the actuators had different constructions, we normalized the performance of a unitary actuator using our ring-like actuator topology as the baseline. Specifically, if a prior actuator had a fiber-like structure, we considered the force and strain generated by two strands of muscle fibers. Alternatively, if the muscle actuator was sheet- or stripe-like, we considered the force generated by the entire unit without stacking. Force and strain data were directly acquired from the literature if they were reported in text. If a performance value was not available in writing or in the supplemental dataset, the value was then extracted from the relevant figures using ImageJ and pixel measurements. The data used to generate Figure 1C are presented in Table S8 and Table S9.

Data processing methods

***Statistical uncertainty propagation:*** Stiffness measurements acquired from the physical compliant skeleton characterization were used to estimate the force output of muscle actuators. To propagate error and uncertainty, we used the formula $\sigma_z = (\bar{x} * \bar{y}) * \sqrt{\frac{\sigma_x}{\bar{x}}^2 + \frac{\sigma_y}{\bar{y}}^2}$ when multiplying the skeleton compression measurements with the characterized skeleton stiffness to find the actuator force output. The $\sigma$ terms were the standard deviation of a measurement, and barred symbols were the mean of the same measurement.

***Images and video processing:*** All photos and videos of devices and actuators were processed and composed with Adobe Photoshop (v26.8.1), Premiere Pro (v25.3), and Illustrator (v29.6.1). No alterations were made to the images used in this work, except for adjusting brightness and contrast for improved visibility. All renders were generated in Rhinoceros 3D version 8.20.

***Myotube alignment analysis:*** The images shown in Figure 3E were processed in the image processing software Fiji to study the alignment of myotubes. The plot shown in Figure 3G were generated from the result of *Directionality* analysis provided by the software.

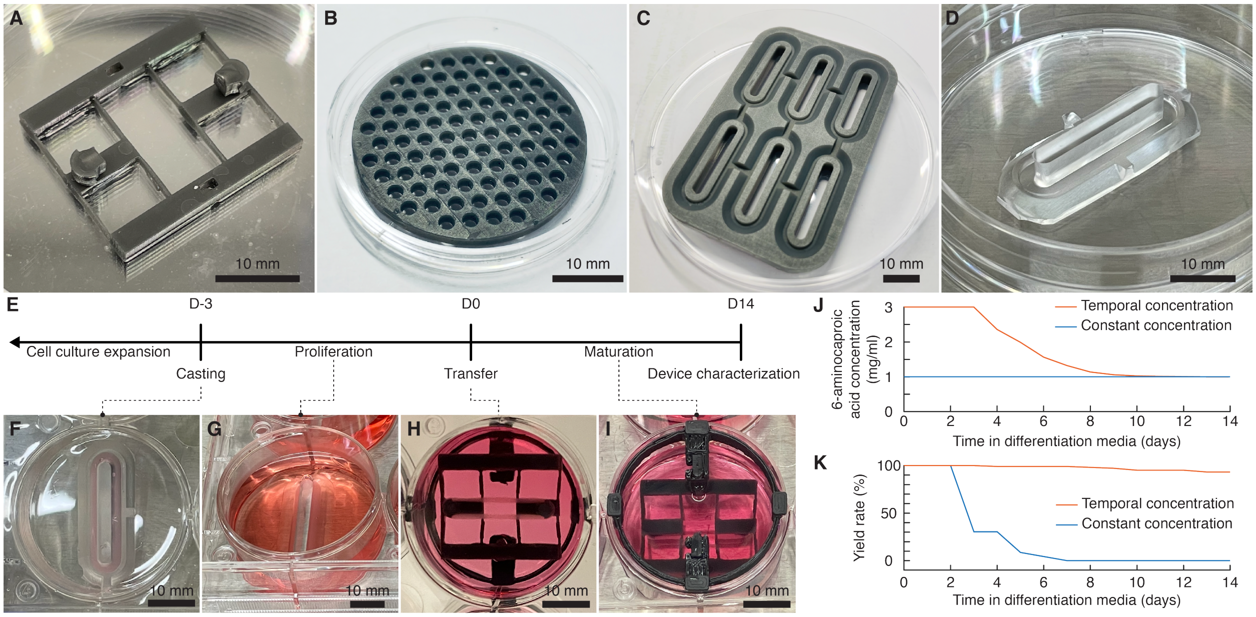


**Figure S1**: **Suspended compliant skeleton and muscle actuator fabrication.** A) Photo of a prepared 20 mm, 1.121 mN/mm compliant skeleton. B) Photo of the magnetic interface (iron-PDMS insert) casting mold. C) Photo of a 20 mm muscle ring master mold. D) Photo of a 20 mm PDMS muscle ring casting mold. E) Muscle actuator fabrication and maturation timeline. F) Photo of a casted, 20 mm muscle ring. G) Photo of a 20 mm muscle ring submerged in supplemented growth media for cell proliferation. H) Photo of an assembled 20 mm, 1.121 mN/mm compliant skeleton and muscle actuator viewed from below. I) Photo of a 20 mm, 1.121 mN/mm compliant skeleton and muscle actuator in supplemented differentiation media for maturation viewed from above. J) Plot of 6-aminocaproic acid concentration versus maturation time. K) Plot of yield rate versus maturation time using different 6-aminocaproic acid concentration schedules. Scale bars, 10 mm.

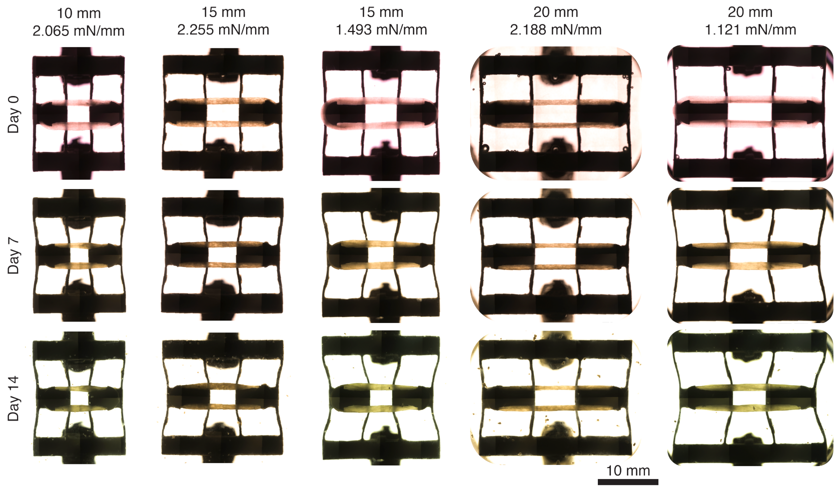


**Figure S2**: **Images of muscle actuator passive compression over time.** Scale bar, 10 mm.

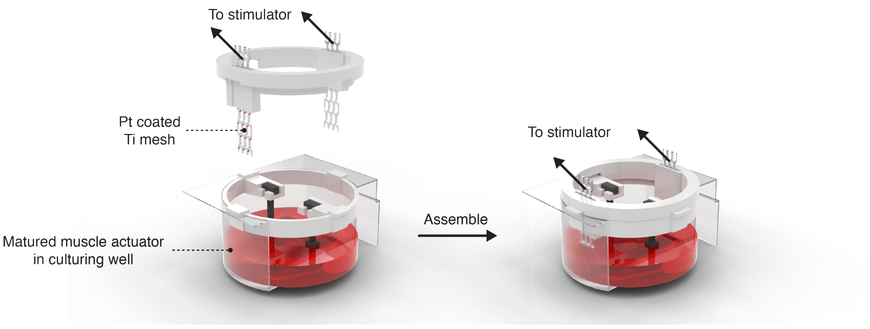


**Figure S3**: **Computer render of the stimulation insert for muscle actuator characterization.**

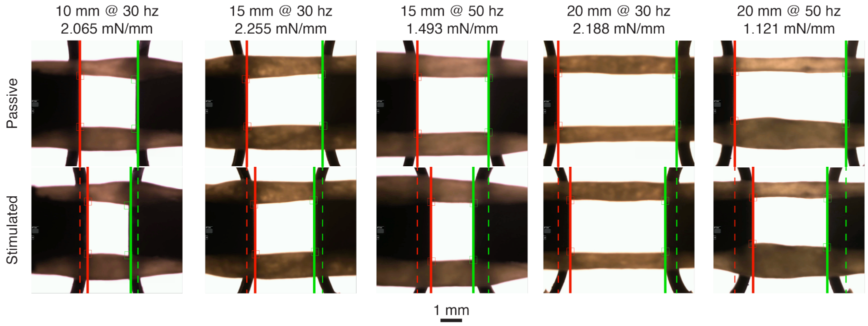


**Figure S4**: **Images of varying muscle actuator and compliant skeleton compression under their peak stimulation frequency.** Scale bar, 1 mm.

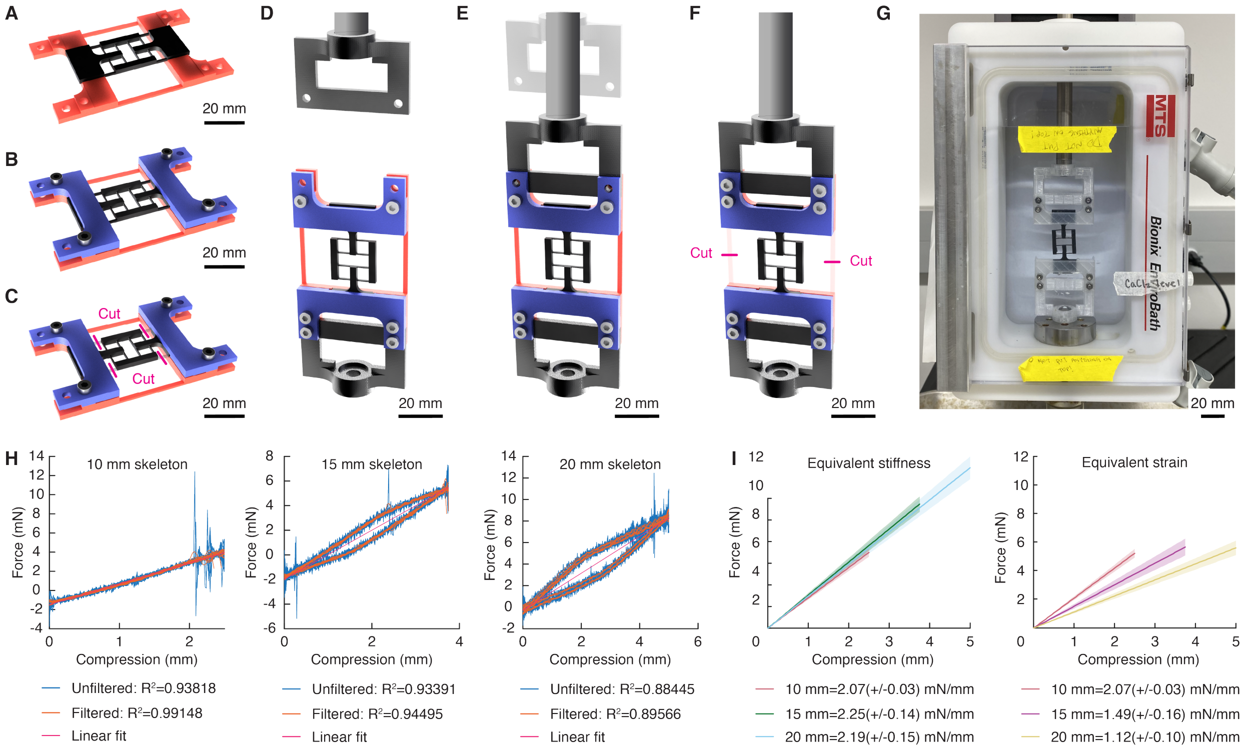


**Figure S5**: **Compliant skeleton characterization methods.** A-C) Assembly of compliant skeleton into the test frame (CAD render). D-F) Assembly of the test frame into the MTS crosshead (CAD render). G) Photo of compliant skeleton test frame assembly in filled MTS biobath prior to testing. Scale bars, 20 mm. H) Filtering and fitting of raw data in MATLAB for buoyancy groups. I) Plotted mean stiffness (Force vs Compression) and standard deviation of each skeleton sample group.

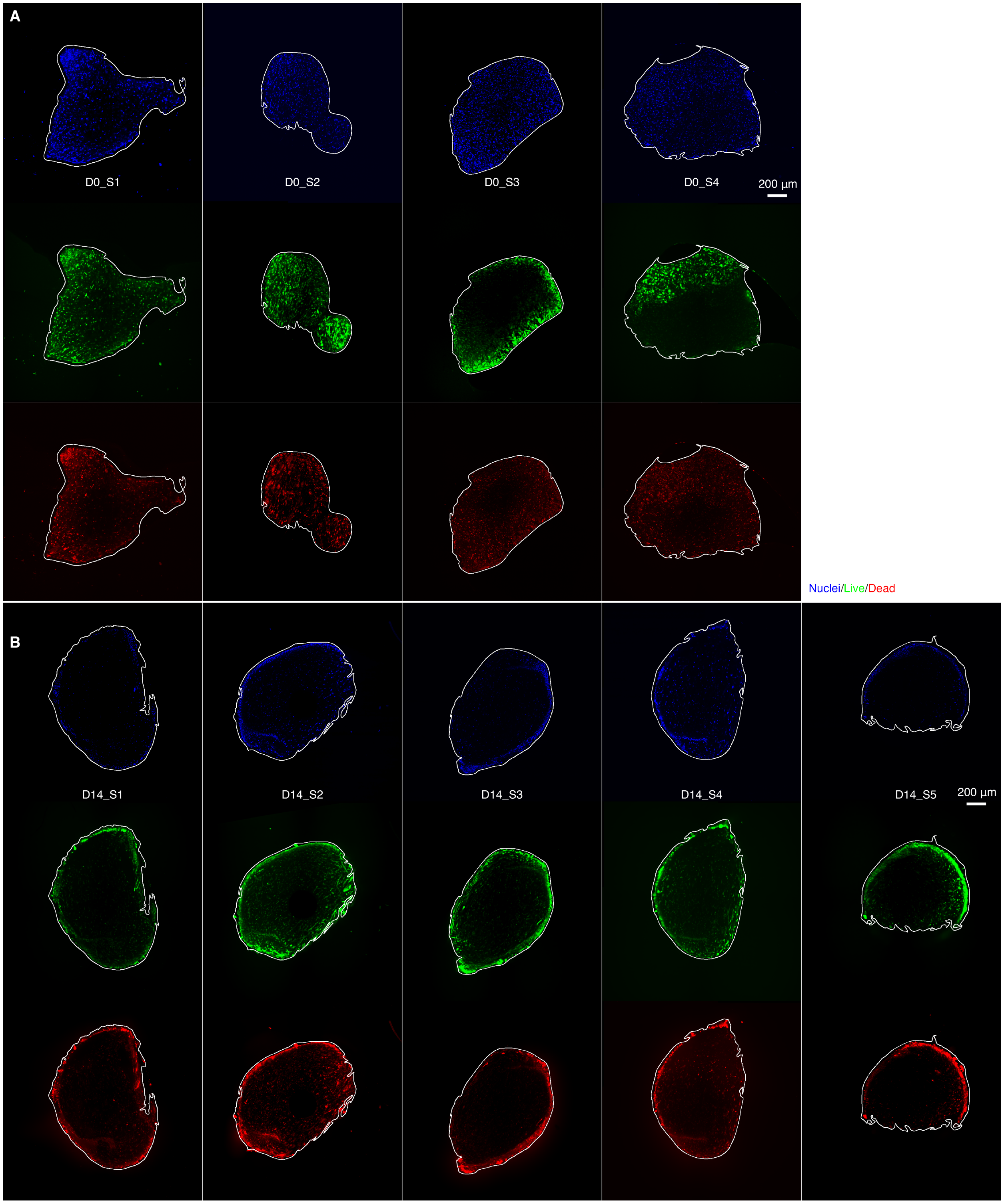


**Figure S6**: **Live/dead cross-section images.** A) D0 cross-section images. B) D14 cross-section images. The contours calculated by the analysis tool were overlaid in white. white outlines show the contours calculated by the analysis tool. Blue, nuclei; green, live cells; red, dead cells. Scale bars, 200 $\mu$m

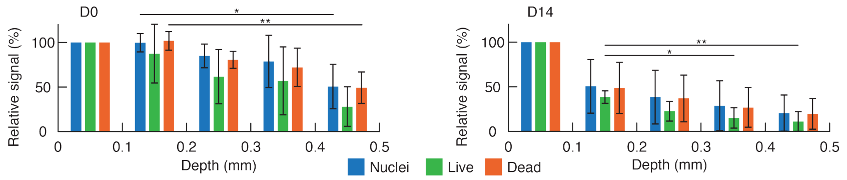


**Figure S7**: **Muscle actuator fiber cross-section live/dead analysis.** All data are mean±SD. D0, n=4; D14, n=5. Statistically significant difference threshold for Tukey's post hoc test: *, $0.01 \leq p < 0.05$; **, $0.001 \leq p < 0.01$.

**Table S1**: **Typical muscle actuator characterization stimulation schedule** This table reports the typical stimulation schedule used to characterize the matured muscle actuators.

| Stimulation count | Start time (s) | Frequency (Hz) |
|---|---|---|
| 1 | 0 | 1 |
| 2 | 30 | 5 |
| 3 | 60 | 10 |
| 4 | 90 | 20 |
| 5 | 120 | 30 |
| 6 | 150 | 50 |
| 7 | 180 | 50 |
| 8 | 210 | 50 |
| 9 | 240 | 50 |
| 10 | 270 | 50 |
| 11 | 300 | 50 |
| 12 | 330 | 50 |
| 13 | 360 | 50 |
| 14 | 390 | 50 |
| 15 | 420 | 50 |

**Table S2**: **Mechanical properties of samples by flexure length.** This table presents the mean calculated stiffness, standard deviation, and relative standard deviation for each sample group of a different length and intended flexure compliance.

| Design | N | Mean Stiffness (mN/mm) | Std | Std % |
|---|---|---|---|---|
| 10 mm, equivalent stiffness and equivalent strain | 3 | 2.0646 | 0.1187 | 5.75 |
| 15 mm, equivalent stiffness | 3 | 2.2549 | 0.1400 | 6.21 |
| 20 mm, equivalent stiffness | 3 | 2.1875 | 0.1540 | 7.04 |
| 15 mm, equivalent strain | 3 | 1.4930 | 0.1570 | 10.51 |
| 20 mm, equivalent strain | 3 | 1.1212 | 0.1001 | 8.93 |

**Table S3**: **$R^2$ values for filtered and fitted compliant skeleton stiffness data.** This table reports the R-squared values for each tested sample pre- and post-filtering with respect to the linear fit equation $y = a * x + b$.

| Design | Sample ID | Unfiltered $R^2$ | Filtered $R^2$ |
|---|---|---|---|
| 10 mm, equivalent stiffness and equivalent strain | 1 | 0.9733 | 0.9841 |
| 10 mm, equivalent stiffness and equivalent strain | 2 | 0.9791 | 0.9891 |
| 10 mm, equivalent stiffness and equivalent strain | 3 | 0.9766 | 0.9890 |
| 15 mm, equivalent stiffness | 1 | 0.9745 | 0.9793 |
| 15 mm, equivalent stiffness | 2 | 0.9769 | 0.9803 |
| 15 mm, equivalent stiffness | 3 | 0.9775 | 0.9811 |
| 20 mm, equivalent stiffness | 1 | 0.9567 | 0.9593 |
| 20 mm, equivalent stiffness | 2 | 0.9633 | 0.9656 |
| 20 mm, equivalent stiffness | 3 | 0.9633 | 0.9659 |
| 15 mm, equivalent strain | 1 | 0.9711 | 0.9737 |
| 15 mm, equivalent strain | 2 | 0.9730 | 0.9761 |
| 15 mm, equivalent strain | 3 | 0.9739 | 0.9773 |
| 20 mm, equivalent strain | 1 | 0.9478 | 0.9522 |
| 20 mm, equivalent strain | 2 | 0.9362 | 0.9403 |
| 20 mm, equivalent strain | 3 | 0.9432 | 0.9480 |

**Table S4**: **Schedule of partial media changes for ACA taper**. This table outlines the media change schedule used to gradually reduce the ACA concentration from the initial 3 mg/mL to 1 mg/mL over the 14-day culture period.

| Day | Media volume in well (ml) | Calculated ACA concentration in well (mg/ml) | Media volume exchanged (ml) | Concentration of ACA in media added (mg/ml) |
|---|---|---|---|---|
| 0 | 7 | 3 | | |
| 1 | 7 | 3 | 3 | 3 |
| 2 | 7 | 3 | 3 | 3 |
| 3 | 7 | 3 | 3 | 3 |
| 4 | 7 | 2.357 | 3 | 1.5 |
| 5 | 7 | 1.990 | 3 | 1.5 |
| 6 | 7 | 1.566 | 3 | 1 |
| 7 | 7 | 1.323 | 3 | 1 |
| 8 | 7 | 1.139 | 4 | 1 |
| 9 | 7 | 1.059 | 4 | 1 |
| 10 | 7 | 1.025 | 4 | 1 |
| 11 | 7 | 1.011 | 4 | 1 |
| 12 | 7 | 1.005 | 4 | 1 |
| 13 | 7 | 1.002 | 4 | 1 |
| 14 | 7 | 1.001 | 4 | 1 |

**Table S5**: **Yield rate of actuators with and without the ACA taper.** This table details the number of actuators remaining unbroken each day of culture in either the ACA taper group or the constant ACA group, along with the calculated manufacturing yield rate.

| Day | Number of unbroken actuators with ACA Taper | Yield Rate with ACA Taper (%) | Number of unbroken actuators with constant ACA | Yield Rate with constant ACA (%) |
|---|---|---|---|---|
| 0 | 98 | 100 | 23 | 100 |
| 1 | 98 | 100 | 23 | 100 |
| 2 | 98 | 100 | 23 | 100 |
| 3 | 98 | 100 | 7 | 30.435 |
| 4 | 98 | 100 | 7 | 30.435 |
| 5 | 98 | 100 | 2 | 8.696 |
| 6 | 98 | 100 | 1 | 4.348 |
| 7 | 98 | 100 | 0 | 0 |
| 8 | 97 | 98.980 | 0 | 0 |
| 9 | 96 | 97.959 | 0 | 0 |
| 10 | 94 | 95.918 | 0 | 0 |
| 11 | 94 | 95.918 | 0 | 0 |
| 12 | 94 | 95.918 | 0 | 0 |
| 13 | 93 | 94.898 | 0 | 0 |
| 14 | 93 | 94.898 | 0 | 0 |

**Table S6**: **Elephant's foot modeling parameters.** This table details the parameters used to model the elephant's foot effect due to overexposure.

| Parameter | definition | value | unit |
|---|---|---|---|
| $h$ | Compliant flexure height | 1.5 | mm |
| $h_{lower}$ | Compliant flexure height with elephant's foot | 0.7 | mm |
| $h_{upper}$ | Compliant flexure height without elephant's foot | 0.8 | mm |
| $b$ | Compliant flexure width | 0.35 | mm |
| $K_{exp}$ | Characterized compliant skeleton stiffness | 2.065 | mN/mm |
| $K_{baseline}$ | Simulated compliant skeleton stiffness (without elephant's foot) | 0.911109 | mN/mm |
| $K_{diff}$ | Difference between characterized and simulated baseline compliant skeleton stiffness | 2.266468 | unit-less |
| $I_{exp}$ | Second moment of area of compliant flexure crossection with the elephant's foot | 0.012146 | $mm^4$ |
| $I_{baseline}$ | Second moment of area of compliant flexure crossection without the elephant's foot | 0.005359 | $mm^4$ |
| $w$ | Elephant's foot widening | 0.192 | mm |
| $o$ | Elephant's foot perimeter offset | 0.096 | mm |

**Table S7**: **Cross-section live/dead analysis processing parameters.** This table details the parameters used by the computational pipeline to generate the contours of cross-section images, which were subsequently used to generate the plots in Figure 3I and contours in Figure S6.

| Cross-section name | Block size | C | Kernel size | Iterations |
|---|---|---|---|---|
| D0_S1 | 21 | 3 | 3 | 0 |
| D0_S2 | 9 | 1 | 3 | 0 |
| D0_S3 | 4 | 1 | 3 | 0 |
| D0_S4 | 7 | 1 | 3 | 0 |
| D14_S1 | 5 | 0 | 0 | 0 |
| D14_S2 | 7 | 3 | 3 | 0 |
| D14_S3 | 5 | 1 | 3 | 0 |
| D14_S4 | 9 | 1 | 3 | 0 |
| D14_S5 | 7 | 0 | 0 | 0 |

**Table S8**: **Literature muscle actuator performance benchmark.** This table details the origins and performances of muscle actuators reported by the literature that were used to produce Figure 1C.

| Reference number | Cell type | Characteristic length (mm) | Active force (mN) | Active strain (%) | Source |
|---|---|---|---|---|---|
| 31 | C2C12 | 5 | 0.3 | 5.662 | Figure 6 |
| 32 | Human skeletal muscle | 40 | 1 | 9.38 | Figure 3 |
| 33 | Harvested murine skeletal muscle | 17.648 | 1.9 | 6.20 | Figure 5 |
| 36 | C2C12 | 15 | 0.162 | 2.50 | Figure 4, Supplemental Material |
| 47 | C2C12 | 4 | 0.3 | 20.00 | Figure 2 |
| 48 | C2C12 | 20 | 0.02 | 4.00 | Figure 3, Figure 6 |
| 49 | C2C12 | 1 | 0.0012 | 0.80 | Figure 5 |
| 50 | C2C12 | 12 | 0.44 | 8.33 | Abstract, Figure 6-9 |
| 51 | C2C12 | 0.5 | 0.00764 | 4.00 | Supplemental Material |
| 52 | C2C12 | 14 | 1.140 | 5.00 | Figure 5 |
| 53 | C2C12 | 6 | 0.5 | 16.67 | Figure 3, Figure 4 |
| 54 | Murine cardiomyocyte | 0.0785 | 0.191 | 11.46 | Figure 5 |
| 55 | Murine cardiomyocyte | 10 | 0.0742 | 3.00 | Figure S7 |

**Table S9**: **Muscle actuator performance benchmark.** This table details the performances of muscle actuators cultured on different compliant skeletons that were used to produce Figure 1C. The characteristic length was set as the length of the muscle actuator.

| Characteristic length (mm) | Skeleton stiffness (mN/mm) | Stimulation electric field frequency (Hz) | Active force (mN) | Active strain (%) |
|---|---|---|---|---|
| 20 | 1.121 | 50 | 2.062 | 10.56 |
| 20 | 1.121 | 30 | 1.580 | 8.09 |
| 20 | 2.188 | 30 | 2.089 | 5.20 |
| 15 | 1.493 | 50 | 1.525 | 8.02 |
| 15 | 1.493 | 30 | 1.413 | 6.70 |
| 15 | 2.255 | 30 | 1.664 | 5.68 |
| 10 | 2.065 | 50 | 1.489 | 8.35 |
| 10 | 2.065 | 30 | 1.194 | 6.70 |

**Movie S1. 10 mm muscle actuator spontaneous contraction.** This video is an animated version of Figure 3B, showing the spontaneous contractions observed on day 7, 10, and 14 with plots to illustrate the patterns produced the 10 mm actuators.

**Movie S2. 15 mm equivalent strain stimulation.** This video is an animated version of Figure 4B, showing the muscle actuator's contraction in response to electrical stimulation (0.353 V/mm, 10 ms pulse width, biphasic) at 1, 5, 10, 20, 30, and 50 Hz.

**Movie S3. Muscle actuator peak stimulation response.** This video is an animated version of Figure 1B and Figure 4A-B, showing the muscle actuators' peak contraction in response to electrical stimulation (0.353 V/mm, 10 ms pulse width, biphasic) at 30 Hz. The muscle actuators were cultured on varying compliant skeleton types. Additional 50 Hz stimulation response is provided for the 15 mm and 20 mm muscle actuators in the equivalent strain group.

**Movie S4. Parallel actuator arrangement.** This video is an animated version of Figure 5E, showing the parallel muscle actuator device's assembly, performance, damage, and repair.

**Movie S5. Biohybrid gripper.** This video is an animated version of Figure 5F-G, showing the biohybrid gripper's operation in response to electrical stimulation.

**Movie S6. Serial actuator arrangement.** This video is an animated version of Figure 6A-C, showing the robotic arm-like device's performance, damage, and repair.

**Movie S7. 2-DOF positioning stage.** This video is an animated version of Figure 6D-F, showing the 2-DOF positioning stage's operation in response to electrical stimulation.

**CAD files.** **Compliant skeletons and accessories.** The following digital models are available in both .stl and .stp format:

- Iron-PDMS insert mold
- 10 mm muscle ring master mold
- 15 mm muscle ring master mold
- 20 mm muscle ring master mold
- 10 mm compliant skeleton
- 15 mm equivalent stiffness compliant skeleton
- 15 mm equivalent strain compliant skeleton
- 20 mm equivalent stiffness compliant skeleton
- 20 mm equivalent strain compliant skeleton
- Equivalent stiffness compliant skeleton hanger
- 15 mm equivalent strain compliant skeleton hanger
- 20 mm equivalent strain compliant skeleton hanger
- Equivalent stiffness compliant skeleton well insert
- 15 mm equivalent strain compliant skeleton well insert
- 20 mm equivalent strain compliant skeleton well insert
- Electrical stimulation insert
